# Subcellular spatiotemporal proteomics delineates distinct phases of ER stress proteostatic response

**DOI:** 10.64898/2026.09.25.754245

**Authors:** Lorena Alamillo, Thalia Paiz, Alexander Black, Matthew C. Juber, Ethan M. Arends, Jacob G. Cox, Dominic C.M. Ng, Timothy A. McKinsey, Maggie P.Y. Lam, Edward Lau

## Abstract

Endoplasmic reticulum (ER) stress is associated with many human diseases, but current understanding of how ER stress responses reshape the stressed proteome over time is still emerging. Specifically, how organellar protein quality control and clearance pathways coordinate to maintain proteostasis in early and prolonged ER stress is unclear. Here we describe a spatiotemporal proteomic strategy termed simultaneous proteome localization and turnover analysis with time resolution (SPLAT-TR) to interrogate the synthesis, degradation, and localization changes of over 4,000 proteins in early (1-4 hours) and prolonged (16-24 hours) ER stress. As ER stress progresses, each time point is distinguished by distinct protein translocation and clearance regulations, coupled to differential organellar proteostasis and usage of protein degradation pathways. Mitochondria show a bimodal response, with a shift from early activation of respiratory protein towards protein quality control pathways in prolonged stress, coinciding with energetics decline. In the Golgi, the increased clearance of collagen proteins is paradoxically coupled to an increased synthesis of secretory pathway components, suggesting removal by secretion. In the ER, the proteome remodels under prolonged stress via increased synthesis of UFMylation and ER-phagy-related proteins coupled to the selective degradation of ER membrane and microdomain proteins. Inhibition of UFMylation alters ER-phagy receptor usage and synergistically induces stress-induced cell death. These results present a systematic delineation of cell compartment proteostasis under unfolded protein response and highlight the utility of SPLAT-TR to investigate time-dependent cellular events.

## INTRODUCTION

ER stress is associated with diseases ranging from cardiovascular pathologies, neurological degradation, cancer, to diabetes (1–3). Under conditions that disrupt protein integrity including calcium imbalance, reactive oxygen species, or genetic mutations, loss of protein homeostasis and folding capacity in the ER triggers the unfolded protein response (UPR). UPR comprises a network of orchestrated cellular events governed by a hierarchical timescale (4). Within minutes of exposure to stressors, three ER-localized receptors IRE1ɑ, PERK, and ATF6, normally bound to BiP/Grp78, are released to activate downstream pathways (5, 6). In the acute-phase response, translational and post-translational regulation halts global protein synthesis within 15–30 minutes (4, 7), which is then followed by transcriptional regulations along the ATF4, IRE1ɑ-XBP1s and ATF6 arms to induce the transcription of stress response genes within 1–2 hours (1, 8, 9).

Following the well-described early signaling events, the downstream molecular sequelae of stress response and their effects on the proteome time evolution remain poorly defined. Although a robust transcriptional activation of UPR target genes can be seen within 1–6 hours (9, 10), changes in proteome composition both poorly correlate with transcriptome changes and lag the transcriptome activation, with the proteome response becoming most robustly observable only at 16 to 24 hours (11). Moreover, only limited overlaps are observed in gene expression changes across timepoints (11), underscoring the continual time evolution and dynamism of stress response programs. Therefore, cellular response to ER stress is marked by a hierarchy of signaling events, broad proteome changes uncoupled from transcript profiles, and still unfilled gaps in proteostatic remodeling that would have implications for understanding disease mechanisms.

Advances in proteomics technologies have led to methods that can reveal proteome dynamics parameters not captured in transcript and protein abundance measurements alone. Stable isotope labeling by amino acids in cell culture (SILAC) approaches allow protein turnover half-life to be measured from their isotope replacement kinetics (12–15). In parallel, subcellular spatial proteomics enables unbiased discovery of protein subcellular localization and differential localization (16, 17). In prior work, we integrated dynamic SILAC and ultracentrifugation-based subcellular proteomics approach to examine the proteome response to ER stressors (18), and observed widespread protein translocation as a previously underappreciated feature of ER stress response. Despite these advances, continued development is needed to resolve global proteostatic status in ER stress response with high spatial and temporal resolution, and the cellular proteome landscape of the ER stress response remains to be fully explored.

Here, we describe a mass spectrometry-based proteomic method, SPLAT-TR, to investigate protein differential localization events, while separately estimating protein synthesis and degradation across subcellular compartments, and their changes across stress response timepoints. Applying SPLAT-TR, we performed spatiotemporal proteomics analysis to assess proteome-wide progression of stress response over distinct timepoints from early to prolonged ER stress. The data reveals an orchestrated protein translocation program that proceeds sequentially between early and prolonged ER stress, and a propagation of stress response to distinct organelles over time. Moreover, our localization-informed clearance and synthesis measurements suggest an underexplored mechanism of collagen removal via increased secretion and implicate a connection between UFMylation and specific ER-phagy receptor usage under prolonged ER stress. Together, these results illuminate the spatiotemporal dynamics of the proteome and expand current view into the molecular changes following ER stress response. The protein synthesis and degradation data at individual protein and organelle resolution reveals new molecular phenotypes and intervention opportunities to target cellular stress response and survival.

## RESULTS

### SPLAT-TR enables protein synthesis and degradation profiling across cellular compartments

Subcellular spatial proteomics presents a powerful method to assign protein subcellular localization and detect localization changes in an unbiased manner. In typical ultracentrifugation-based workflows, a gentle cell lysis containing unruptured organelles are centrifuged over a density gradient or at differential speeds to create centrifugation fractions (19–22). Following tandem mass tag (TMT) labeling and mass spectrometry quantification of protein fractionation profiles, a semi-supervised model is trained on the fractionation profile using a list of compartment marker proteins to reconstruct protein spatial distributions. The model then classifies each of the unknown proteins into a subcellular designation depending on its distributional pattern. In recent work, we devised a spatiotemporal proteomics workflow termed simultaneous protein localization and turnover analysis (SPLAT) (18, 23), which builds on the ultracentrifugation-based subcellular fractionation protocol LOPIT-DC (20) as its spatial enrichment engine, and additionally incorporates dynamic stable isotope labeling of amino acid in cell culture (dynamic SILAC) to simultaneously measure protein translocation and turnover rates within the same cells. SPLAT allows spatially resolved measurements of protein turnover kinetics and inference of protein trafficking defects based on the differential localization of newly synthesized proteins. However, a major limitation of SPLAT is that it does not separately estimate the respective contributions of protein synthesis and degradation to protein turnover under non-equilibrium conditions. Moreover, the protocol requires cellular content to be separated into 10 ultracentrifugation fractions to achieve subcellular spatial resolution, thus limiting throughput and sensitivity and restricting prior comparisons to single timepoints.

To further characterize the global proteostasis changes during progressive stress response, we describe the SPLAT with time resolution (SPLAT-TR) method. SPLAT-TR re-designs the SPLAT workflow to enable separate measurements of protein synthesis and clearance while increasing throughput to interrogate multiple stress response timepoints or treatment conditions (**Figure 1**). In typical dynamic SILAC experiments, the reported turnover values are calculated from an aggregate of SILAC light (L) and heavy (H) peak signals. To disentangle the L (representing clearance) and H (largely representing synthesis) intensities, a TMT-SILAC “hyperplexing” strategy can be employed (24, 25) where SILAC L and H intensities of individual samples are compared across TMT channels within the same MS2 spectrum. This foregoes MS1-level SILAC-L and SILAC-H normalization and separates synthesis and clearance kinetics of proteins from their composite turnover rates, allowing them to be fairly compared across timepoints and conditions. However, in the LOPIT-DC/SPLAT design, MS2-level multiplexing, i.e., TMT channels, is already used to barcode subcellular localization information by multiplexing sequential ultracentrifugation fractions. To overcome this challenge, we aimed to encode control and treated samples within the same TMT block using available channels, i.e., to encompass spatial information as well as relative quantitative across multiple treatment conditions and/or time-points.

**Figure 1.**
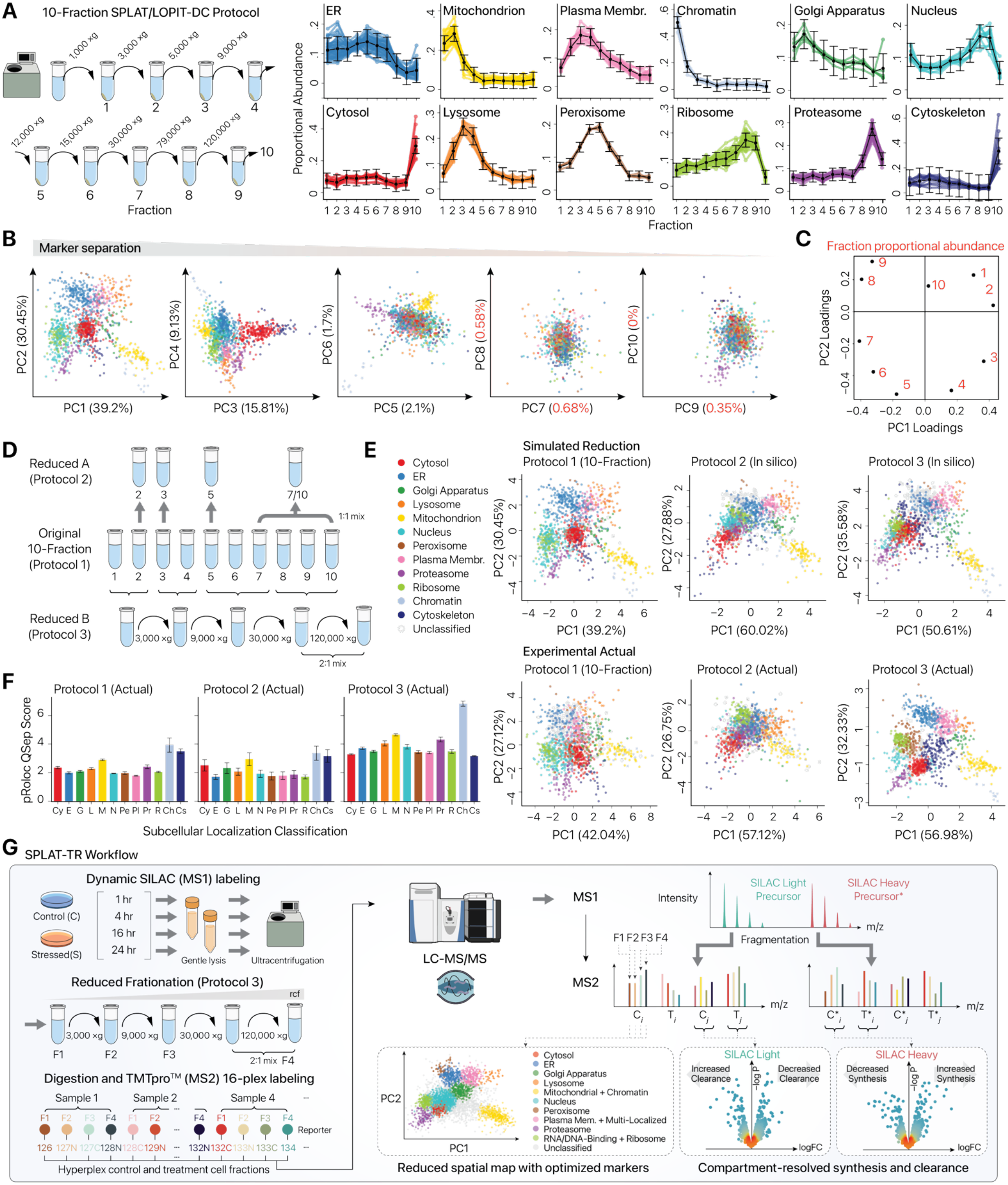
SPLAT-TR enable subcellular localization and synthesis-clearance measurements. (A) Top: Schematic depicting the standard 10-fraction subcellular fractionation protocol (termed Protocol 1) derived from LOPIT-DC. Right: Distribution patterns of normalized abundance (y-axis) calculated from TMT channel intensities for each subcellular marker over the ten ultracentrifugation fractions (x-axis) from the AC16 normal condition in the previous publication on SPLAT. Black lines depict the average intensity at each fraction for the markers of that compartment with error bars for the standard deviation. (B) PCA plots for each of 10 principal components (PC) in SPLAT AC16 normal condition. Each data point is a protein with color corresponding to BANDLE localization classification. (C) Scatter plot of the loadings PC1 (x-axis) and PC2 (y-axis) from AC16 normal condition. (D) Schematic depicting two methods to reduce the fractions from the original 10-fraction SPLAT, here referred to as Protocol 2 and Protocol 3. (E) PCA plots for PC1 (x-axis) and PC2 (y-axis) for one of the three replicates in each fractionation protocol. Top left to right: prior data on SPLAT AC16 normal condition using the standard Protocol 1; in silico projection of Protocol 2 using existing data from SPLAT AC16 normal condition, in silico projection of Protocol 3 using existing data from SPLAT AC16 normal condition. Bottom left to right: new experimental validation data using the standard 10-fraction separation (Protocol 1), experimental execution of fraction reduction method (Protocol 2), experimental execution of fraction reduction method (Protocol 3). All experiments were performed on untreated and non-SILAC-labeled AC16 cells. (F) Bar plots of the mean QSeps score from the experimental execution of the original Protocol 1 with the new dimension-reduced protocols. Values are calculated from the average of the ratios of each distance between a given cluster and each other cluster over the dispersion of the given cluster. Error bars are standard deviation between replicates (n=3). (G) Overall schematic of the SPLAT-TR method, depicting the use Protocol 3 combined with SILAC on a time course of thapsigargin exposure. Cells are exposed to heavy labeled arginine and lysine in control and thapsigargin-treated conditions. Timepoints begin with the exposure to both heavy and light SILAC and beginning of treatment conditions. The cells are harvested and differentially centrifugated into a series of fractions. This is followed by further processing and TMT labeling to combine two conditions for two timepoints to directly compare synthesis (SILAC heavy) and clearance (SILAC light) changes across ER stress timepoints. The markers and cluster assignments were optimized for the data collected resulting in the portrayed compartments.

To increase multiplexing capability while retaining spatial resolution, we reason we could reduce the number of centrifugation fractions and hence TMT channels needed to represent protein spatial information. In the standard LOPIT-DC/SPLAT protocols, 10 TMT channels are used to represent 10 fractions from 9 ultracentrifugation steps (**Figure 1A**). Upon analyzing existing data using PCA-based spatial maps however, we find that most spatial separation of proteins is contained within the first 4 principal components despite there being up to 10 input spatial dimensions (subcellular fractions/TMT channels) (**Figure 1B**). This suggests fewer number of centrifugation fractions may be sufficient to distinguish cellular compartments effectively.

To compress the ten-step centrifugation protocol we previously employed (“**Protocol 1**”), we examined the loadings of PC1 and PC2 (**Figure 1C**), which represent the contribution of each of the original variables (centrifugation fractions) to the top two PCs, and designed two reduced-fraction protocols (“**Protocol 2**” and “**Protocol 3**”) (**Figure 1D; Supplementary Table S1**). In Protocol 2, we selected individual fractions from the original SPLAT protocol to be included based on their contribution to overall data variance and the organelle proteins that they represent. We included Fractions 7 and 2 as they have the highest contribution to PC1, Fraction 10 because it contains most of the cytosolic proteins, and Fractions 3 and 5 because they have a high contribution to PC1/PC2, respectively, while covering proteins from various organelles. To make a total of four fractions, we combined Fraction 10 with Fraction 7 at equal mass ratios.

In Protocol 3, fractions that cluster closer together are interpreted to contain similar spatial information. As the protein concentrations in the fractions to be combined were roughly equal, we combined each quartile of the PCA plot into one individual centrifugation step to a total of four ultracentrifugation steps. Protocol 1’s Fraction 10, which contains the soluble proteins in the final supernatant, is extracted via acetone precipitation and combined at a 1:2 mass ratio with the single centrifugation step that encompasses Fractions 8 and 9.

To evaluate the spatial resolution afforded by Protocols 2 and 3, we first computationally simulated their subcellular compartment separation by subsetting and combining the fractional data generated for our previous work (**Figure 1E** top). After qualitatively ensuring the two proposed methods lead to clustering and separation between the proteins of the organelles examined, we experimentally compared all three protocols (**Figure 1E** bottom). To keep the comparison consistent, only the proteins present in all data sets (simulated and experimental) were analyzed. Qualitatively, Protocol 3 appeared to show the best clustering and separation compared to both Protocol 2 and the original ten-fraction SPLAT protocol. To quantitatively compare spatial separation, we calculated the Qsep distance (20) to measure the resolution of a spatial proteomic experiment by calculating the distance between each organelle cluster over each within cluster distance (see **Methods**). Intriguingly, Protocol 3 not only achieves better spatial resolution than Protocol 2 but also the ten-fraction Protocol 1 (**Figure 1F**). This runs counter to the intuition that more fractionation steps would enhance spatial resolution, which we speculate may be due to the increased signal-to-noise ratios from combining similar fractions and reduced technical errors from manual pipetting of supernatant from the pellets after each spin. The protein fraction distribution patterns of each protocol likewise show Protocol 3 affords more distinct distribution patterns for ER, Golgi, and the plasma membrane containing fractions (**Supplementary Figure S1**), resulting in cleaner separation on the PCA spatial map (**Figure 1E**). As an additional merit, Protocol 3 also led to more protein identification.

We therefore proceeded with Protocol 3 to resolve subcellular distribution of proteins at 1, 4, 16, 24 hours upon the introduction of ER stress by 1 µM thapsigargin (**Figure 1G**). The timepoints were chosen to represent both early ER stress (1 and 4 hours) and cellular responses to prolonged ER stress (16, 24 hours). By introducing SILAC L/H arginine and lysine concomitant to stressors, this design allows for multiplexing of two timepoints and two conditions into one 16-plex TMT labeling experiment. To reduce potential bias across TMT blocks, we multiplexed control and thapsigargin samples from one early and one late timepoint together within each TMT experiment. In total, we quantified 4344 proteins across four ER stress and control timepoints. We then created an optimized set of compartment markers for Protocol 3 that led to well-separated distribution profiles for 10 curated subcellular compartments: cytosol, ER, Golgi, lysosome, mitochondria/chromatin, nucleus, peroxisome, plasma membrane/multi-localized, proteasome, and ribosome/RNA/DNA-binding (**Supplementary Figure S2A; Supplementary Data S1**). Proteins assigned to each cluster are enriched with corresponding subcellular compartments and organellar terms, supporting that we successfully incorporated subcellular information encoding into a TMT-SILAC design (**Supplementary Figure S2B**).

### Early and late ER stress cause distinct cell-wide protein spatial remodeling events

We next analyzed the SPLAT-TR data to determine the changing spatial contexts of proteins at various stages of stress response, by using the BANDLE (26) to calculate the probability of protein differential localization and identified confident localization changes between normal and stressed cells at 1, 4, 16, and 24 hours post-stress induction (**Figure 2A**; **Supplementary Data S2**). The overall distribution patterns of subcellular compartments were consistent across timepoints and treatments, suggesting the four-fraction Protocol 3 led to consistent spatial separation in normal and stressed cells (**Figure 2A**). We then focused on the subset of proteins with confident predictions of differential localization, as supported by their differential centrifugation profiles across conditions (**Supplementary Figure S3**).

**Figure 2.**
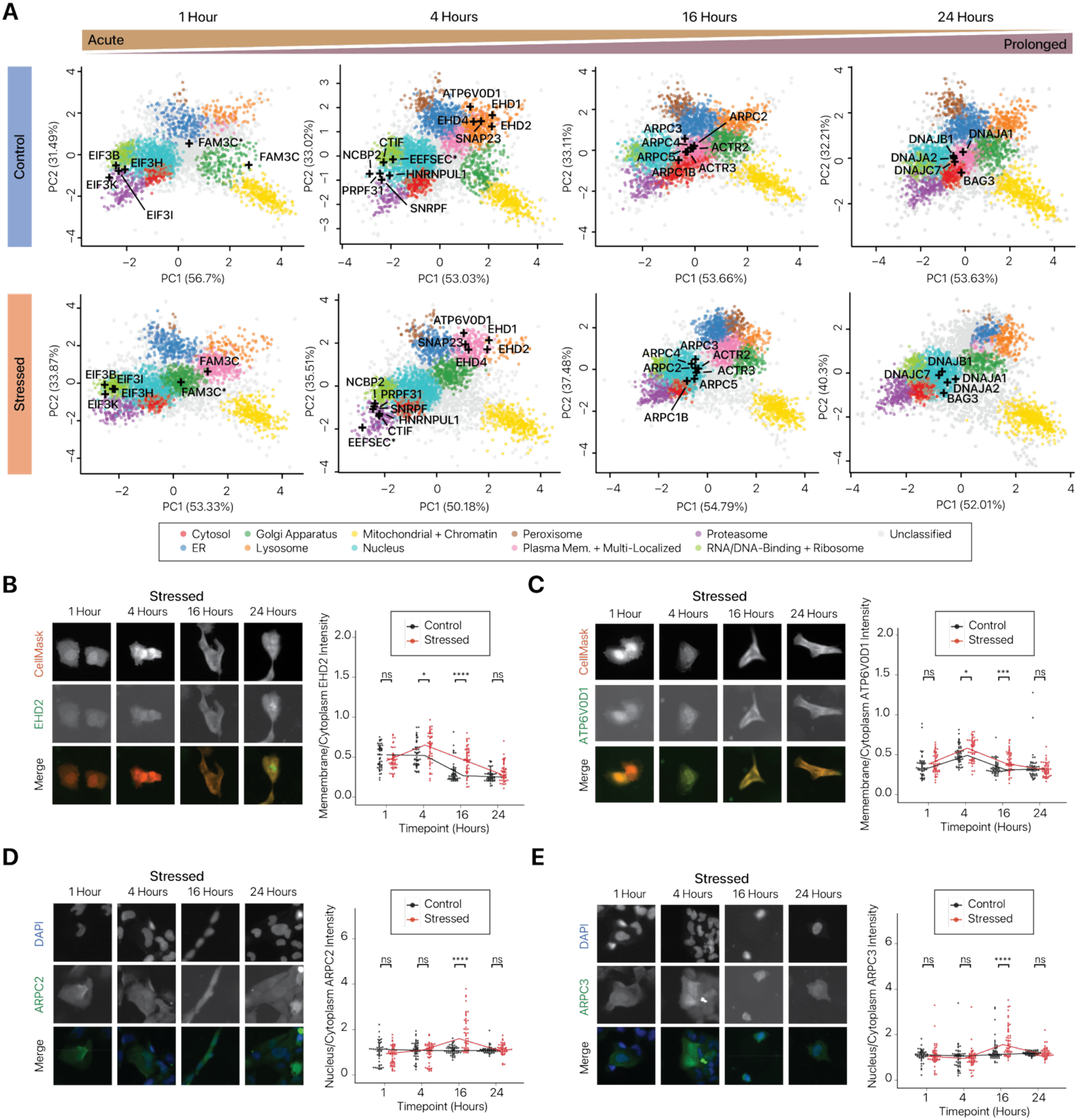
ER stress causes sequential spatial remodeling propagating to neighboring organelles. (A) Spatial maps (PC1/PC2) of normal and stressed AC16 cells at each time point with the subcellular localization (black cross) of highlighted re-localization events at each timepoint. Top panels show the control localizations and bottom panels show the thapsigargin treated localizations. PCAs portrayed are an average of the three replicates collected. Color: subcellular localization assignment; n=3 replicates per timepoint per condition. Asterisks after protein names denote SILAC heavy proteins. (B) Left: High-content imaging of SNAP-tagged constructs for EHD2 in stressed AC16 cells at 1 hour, 4 hours, 16 hours, and 24 hours. Deep read: CellMask for whole cell label; Oregon Green: SNAP ligand. Right: Analysis of the intensity of the Oregon Green SNAP ligand at the membrane over the intensity in the cytoplasm (y-axis) at each timepoint (x-axis). Each timepoint contains 50 cells screened for the highest expression for the protein of interest of the cells imaged at that timepoint. A Shapiro-Wilk test indicated the data deviated from normality. Significance tests were performed using a Wilcoxon rank sum test comparing the control and thapsigargin treated at each timepoint. Solid line indicates the median of each condition. (C) As in B, but for ATP6V0D1. (D) Left: High-content imaging of SNAP-tagged constructs for ARPC2 in stressed AC16 cells at 1 hour, 4 hours, 16 hours, and 24 hours. Blue: nucleus (DAPI); Oregon Green: SNAP ligand. Right: Analysis of the intensity of the Oregon Green SNAP ligand at the nucleus (overlapping DAPI) over the intensity in the cytoplasm (y-axis) at each timepoint (x-axis). Each timepoint contains 50 cells screened for most average nuclear size of the cells imaged at that timepoint. A Shapiro-Wilk test indicated the data deviated from normality. Significance tests were performed using a Wilcoxon rank sum test comparing the control and thapsigargin treated at each timepoint. Solid line indicates the median of each condition. (E) As in D, but for ARPC3.

At 1 hour of ER stress, subunits for eukaryotic initiation factor 3 (EIF3) complex including EIF3B, EIF3I, EIF3H, and EIF3K become differentially localized toward the ribosome-containing RNA-/DNA-binding protein rich compartment. These early events are consistent with stalled ribosome complexes and stress granule formation (27), and EIF3-dependent ATF4 translation (28), as expected from the known stress dependent global translational pause and EIF3-dependent translation during the integrated stress response. In another early event, we highlight that the SILAC-H labeled (newly synthesized) FAM3C differentially localizes from the ER to the Golgi and SILAC-L (pre-existing) FAM3C moved from the Golgi toward the cell surface. FAM3C is known to be secreted during ischemic damage and myocardial infarction (29) whereas our data suggests FAM3C secretion may be a conserved event in early stress response.

At 4 hours of ER stress, multiple mRNA processing enzymes (NCBP2, CTIF, EEFSEC, HNRNPUL1, SNRPF, and PRPF31) move from the nucleus compartment to cytosolic complexes, consistent with continued stress granule recruitment. In other notable events, we highlight multiple endocytosis pathway proteins (ATP6V0D1, SNAP23, EHD1, EHD2, and EHD4) that are predicted to the lysosome compartment in the control condition but are instead predicted to the plasma membrane/multi-localized compartment in stressed cells. To validate the proteomics data, we expressed SNAP-tagged EHD2, EHD4, and ATP6V0D1 in normal and stressed AC16 cells for high-content fluorescent imaging and automated intracellular vs. cell surface fluorescence intensity analysis (**Figure 2B–C; Supplementary Figure S4A**). Corroborating the spatial proteomics data, high-content imaging supports a significant differential localization for EHD2, a membrane/caveolae binding protein involved in endocytosis (**Figure 2B**), and ATP6V0D1, a subunit of a membrane associated proton transport complex (**Figure 2C**). Both proteins show significantly higher membrane-to-cytoplasmic intensity ratio compared to vehicle control at 4 and 16 hours of ER stress but not 1 hour, suggesting their differential localization occurs after acute-phase events. On the other hand, EHD4, an ATP and membrane binding protein also involved in endocytosis, only shows differential localization at 16 and 24 hours (**Supplementary Figure S4A**); we speculate this lag from the proteomics data may be due to detection sensitivity or cell-to-cell variations in the imaging experiment.

At 16 hours, a prominent differential localization involves subunits of the actin-related protein 2/3 (ARP2/3) complex, a cytoskeleton regulator with best characterized role in the cytoplasm where it regulates actin polymerization. ARPC1B, ARPC2, ARPC3, ARPC4, ARPC6, ACTR2, and ACTR3 are classified to become differentially localized toward the nucleus. To validate the proteomics results, we expressed SNAP-tagged ARPC2, ARPC3, and ARPC5 in normal and stressed AC16 cells. The automated high-content imaging analysis confirmed a significant increase in nucleus-to-cytoplasm intensity ratios at 16 hours of thapsigargin treatment compared to vehicle control, in agreement with the proteomics data (**Figure 2D–E; Supplementary Figure S4B**). The differential localization likewise shows an intricate time dependence, with no significant differences at 1 and 4 hours. At 24 hours, ARPC2 and ARPC3 return to baseline while the nucleus-to-cytoplasm intensity ratio for ARPC5 remains significantly elevated. Furthermore, the imaging suggests ARP2/3 is dually localized to the cytoplasm and nucleus at baseline, supporting the allocation of the subunits to the “plasma membrane/multi-localized” compartment in the control condition. The ARP2/3 complex has several described functions in the nucleus associated with nuclear actin, including DNA damage repair (30, 31) and transcription regulation (32). We did not detect the presence of DNA damage over a time course of thapsigargin treatment by immunoblots again γH2AX, a known marker of DNA double stranded breaks (data not shown). Hence, the functional significance of the nuclear shuttling of ARP2/3 remains to be investigated.

Finally, at 24 hours, we highlight several co-chaperones that move from the plasma membrane/multi-localized compartment to the nucleus compartment, including DNAJA1, DNAJB1, DNAJA2, and DNAJC7 (**Figure 2A**, right). This suggests propagation of stress to the nucleus requiring increased protein folding activity. These J-domain protein (JDP) co-chaperones interact extensively with the 70-kDa heat shock protein (HSP70) chaperone network, which shuttles to the nucleus in response to stress (33, 34). Taken together, these results show that SPLAT-TR can resolve spatial dynamics at different timepoints, recapitulating known events (EIF3, stress granules) and nominating time-dependent differential localization events (EHD1/2/4, ARP2/3, and JDP) that can be validated by orthogonal observations. Moreover, the data suggests that as ER stress progresses, there is a sequential spatial remodeling of proteins that propagates to cell-wide organelles at a defined time scale.

### Differential protein clearance and synthesis across organelles under ER stress

To further examine stress propagation to different organelles and how protein synthesis and degradation in each location rates alter, we compared protein synthesis and clearance within the mitochondria, ER, and Golgi apparatus between normal and stressed cells (**Supplementary Data S3**).

For proteins localized to the mitochondrial/chromatin compartment, we note that, at 1 hour, there is an increased synthesis of UQCRC1 and decreased clearance of MT-CO2, ATP5MG, ATP5F1D, ATP5F1B, NDUFV1, and UQCRC1, indicative of an up-regulation of bioenergetics proteins (**Figure 3A**). Notably, these changes are no longer observable at 4 hours of ER stress, and regulation eventually transitions towards an increased synthesis and decreased clearance of key mitochondrial proteostasis machineries at 16 and 24 hours, including HSPD1, LONP1, AIFM1, HSPA9/GRP75, and TOMM70. To evaluate whether this time-dependent alteration of respiratory chain related proteins reflects a sequential alteration of bioenergetics, we performed Seahorse respirometry assays on AC16 cells under early and prolonged stress (**Figure 3B**). The results corroborate a biphasic behavior of mitochondrial respiration, with significant increases in basal respiration, ATP production, and maximal respiratory capacity within 30 minutes of thapsigargin treatment compared to vehicle control. At the same time, however, there is also an increase in proton leakage and decrease in coupling efficiency. By 4 hours, increases in respiration are no longer observed while proton leakage continues to be significantly increased, and coupling efficiency and spare respiratory capacity continue to be significantly decreased. By 24 hours, a significant decrease in basal respiration, ATP-linked respiration, maximal respiration, and spare capacity is observed, corroborating that ER stress affects mitochondrial energetics in an exposure time-dependent fashion (**Figure 3C**).

**Figure 3.**
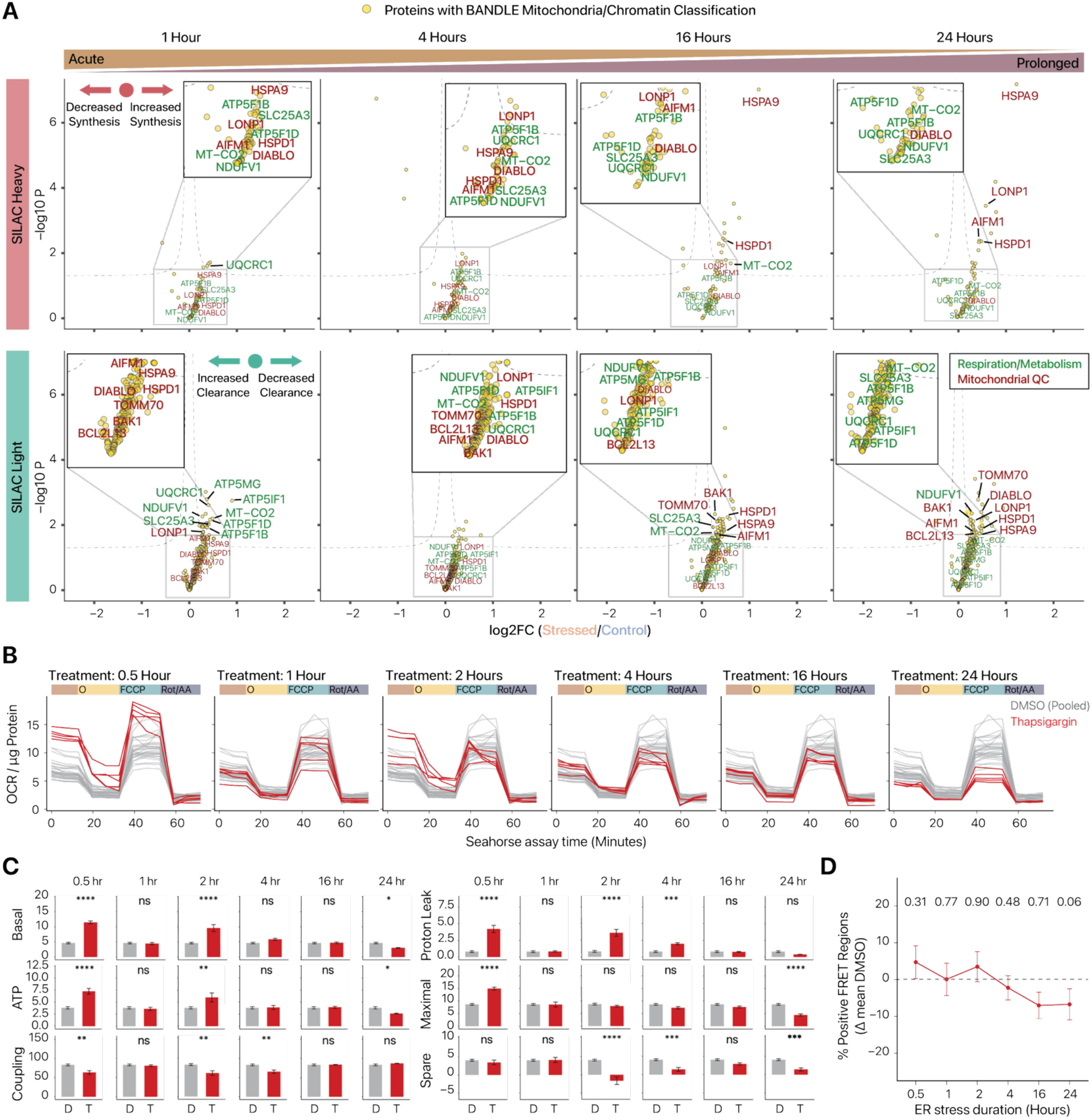
ER stress leads to biphasic changes in mitochondrial energetics proteostasis. (A) Volcano plots of limma contrast fits for the SILAC heavy labeled vs. SILAC light proteins across timepoints in normal and stressed cells. Proteins predicted to reside in the mitochondria/chromatin compartment in 24-hour control cells are included. Y-axis: –log10 of limma p values; x-axis: log foldchange in intensity. The top set of volcano plots is for the heavy labeled proteins from the 1-hour timepoint (left) to the 24-hour timepoint (right). The bottom set of volcano plots is for the light labeled proteins from the 1-hour timepoint (left) to the 24-hour timepoint (right). The proteins annotated in green are proteins associated with energy production. The proteins annotated in red are mitochondrial stress marker proteins. (B) Seahorse assay results with thapsigargin conditions in red and DMSO vehicle control in grey. Oxygen consumption rate normalized for µg of protein per well (x-axis) over the course of the assay (y-axis). Each plot portrays a different timepoint with thapsigargin treated wells represented in red and vehicle control wells in grey. Each condition/timepoint was performed in five biological replicates with all vehicle control replicates across timepoints portrayed on each plot. (C) Summary of Seahorse results. Treatment timepoints are shown from left to right, and assay results are shown from top to bottom. Each condition/timepoint was performed with five biological replicates, each with three technical replicates. The mean of the technical replicates was taken to represent the value of each biological replicate, and the mean of the biological replicates is portrayed by each thapsigargin bar. Vehicle control bars represent controls across all timepoints grouped together. Error bars are the standard error of the mean, and significance was determined using a two-way ANOVA assessing the effects of treatment and time followed by pairwise comparisons with Holm adjustment. (D) Plot for FRET assay with the difference between the mean FRET positive regions per cell in the thapsigargin treated condition to the mean of the FRET positive regions per cell in the DMSO treated condition at each timepoint (y-axis) over the time course (x-axis). Error bars are the standard error of the difference between the two means. Statistics were performed using a Wilcoxon rank sum test between the difference to the mean of the DMSO condition at each timepoint for the DMSO condition and thapsigargin condition.

To further explore the link between early and prolonged ER stress on mitochondrial energetics, we investigated ER-mitochondrial coupling at mitochondria-associated membranes (MAM) or mitochondria-ER contact sites (MERCS), which are known to play a role in ER-mitochondrial crosstalk, calcium transport, and ATP production in early ER stress (35). To measure ER-mitochondrial contact, we expressed a Förster resonance energy transfer (FRET) system of two MERCS components, with the donor on ER resident SEC61B and the acceptor on the mitochondrial membrane protein TOM20 (36). The FRET data corroborate that the number of MERCS as measured via FRET positive regions shows an increase trend in in early ER stress, which then reverts into a trending repression (P = 0.06) at 24 hours (**Figure 3D**). Taken together, the protein synthesis and clearance data, supported with functional assays, portrays a biphasic mitochondrial proteostatic response to ER stress, where the early increase in bioenergetics protein synthesis underlies the increased energy production during the adaptive phase of ER stress response. However, this response appears to be unsustainable as evidenced by the increased proton leak even within 30 minutes of ER stress onset, and gradually leads into depleted energy production whereas the mitochondrial stress response switches to the production of chaperones and proteases to maintain proteostasis.

Cellular proteins are primarily removed through the ubiquitin-proteasome system or autophagy-lysosome pathways, with secretory pathway and membrane-bound organelle proteins preferentially degraded via the latter. To examine the respective contributions of each pathway on the differential protein clearance over time, we examined whether the spatial proteomics data revealed signs of differential protein degradation pathway usages as ER stress progresses. The spatial proteomics data show multiple proteins that differentially localize to compartments associated with protein degradation (**Figure 4A**). We note that such protein compartment changes are dominated by movement toward with proteasome-containing compartment at 1, 4, and 16 hours of ER-stress. Although this does not directly implicate proteasome-mediated degradation, it poses a stark contrast with the prevalent differential localization events towards the lysosome at the 24-hour mark. Moreover, proteins that relocate toward degradation compartments in each timepoint are marked by different functional classes. At 1 hour, multiple RNA-binding proteins shift towards the proteasome; at 4 hours, multiple transcriptional repressors from the nucleus to the proteasome; at 16 hours, multiple cytosolic enzymes to the proteasome; at 24 hours, multiple secretory pathway resident proteins re-localize from the ER and Golgi toward the lysosome. To corroborate this shift, we used an LC3 autophagy flux assay to map the cellular autophagy activity and measured total proteasome capacity to degrade an extrinsic substrate over ER stress timepoints (**Figure 4B–C**). Autophagy activity significantly increases upon the initiation of ER stress and stays consistently upregulated over prolonged ER stress; whereas proteasomal capacity remains constant until the final two timepoints collected where we observed a significant decrease; together suggesting there is a significant shift in utilization of degradation strategies over time as stress response progresses.

**Figure 4.**
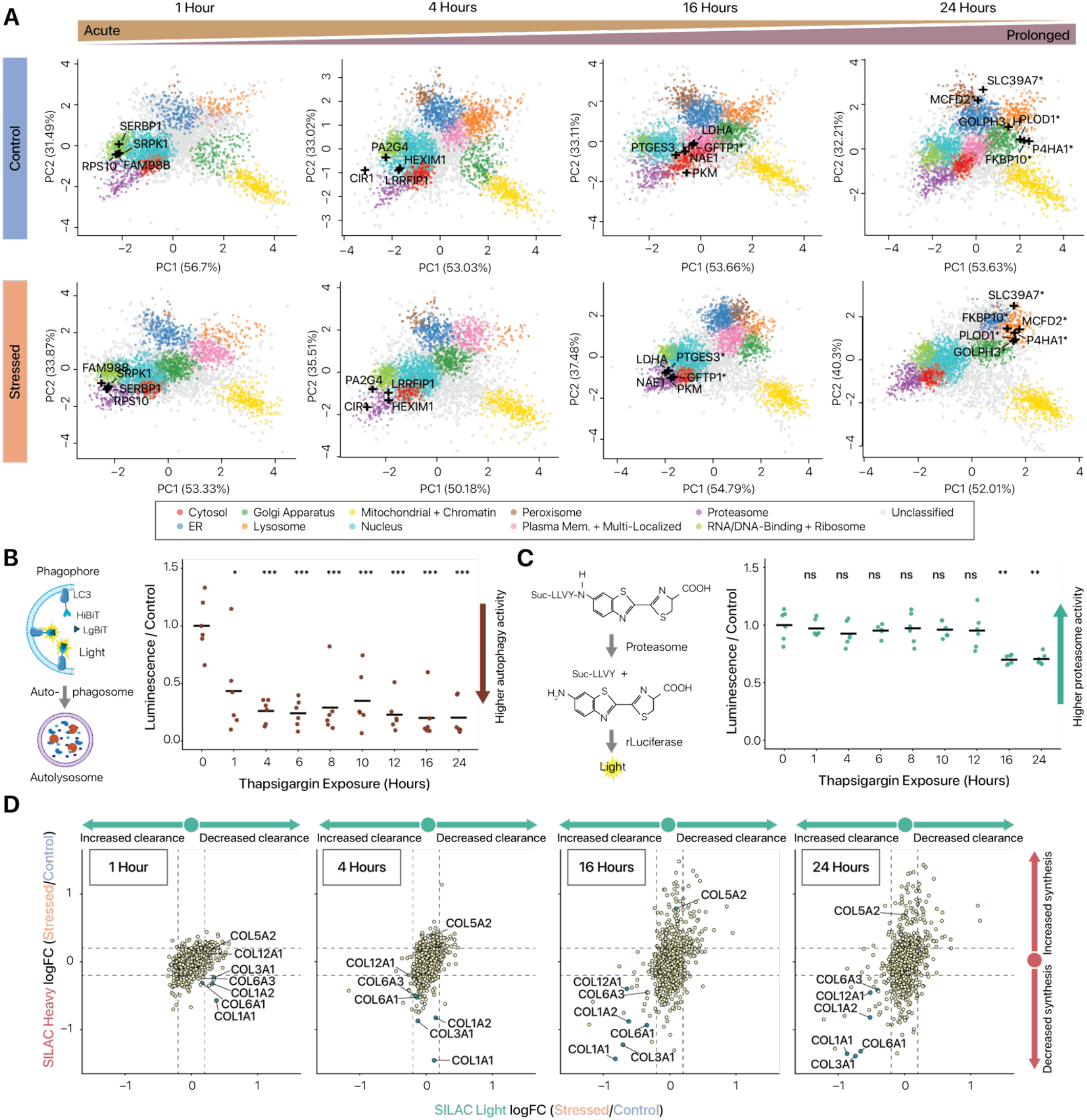
Progressing ER stress induces the clearance of protein targets through multiple protein clearance pathways. (A) Spatial maps (PC1/PC2) of normal and stressed AC16 cells at each time point with the subcellular localization (black cross) of protein predicted to move towards degradative pathways over the time course of ER stress. Top panels show the control localizations, and bottom panels show the thapsigargin treated localizations. PCAs portrayed are an average of the three replicates collected. Legend for color scheme is shown on the right. Asterisks after protein names denote SILAC heavy proteins. (B) Autophagic flux assay over a time course of 1 µM thapsigargin exposure with a schematic portraying assay readout. Increased autophagic flux results in decreased luminescence. Data points are plotted as a ratio to the average of the untreated 0 timepoint. Significance was determined using a t-test comparing each timepoint to the untreated 0 timepoint. Each condition was performed with eight replicates. Solid line indicates the mean of the timepoint. (C) Proteasomal degradative capacity assay over a time course of 1 µM thapsigargin exposure with a schematic portraying assay read out. Increased proteasomal capacity results in increased luminescence and vice versa. Data points are plotted as a ratio to the average of the untreated 0 timepoint. Significance was determined using a t-test comparing each timepoint to the untreated 0 timepoint. (D) Log2 fold change in heavy labeled protein (y-axis) against log2 fold change in light proteins (x-axis) between the 1 µM thapsigargin treated and control at each timepoint. Arrows depict changes in synthesis and clearance. Dashed lines are set at 0.2 log2 fold change.

Interestingly, this decrease in protein degradation capacity at 16 and 24 hours coincides with the stress-induced increase in the clearance rate of a number of proteins, with the strongest signal mapped to collagen proteins, which have been shown to be unamenable to ER-assisted degradation (ERAD) and to require autophagic degradation (37–39). In our data, collagens show increased clearance rates only at prolonged ER stress (**Figure 4D**).

As collagen chains are localized to the Golgi in our experiments in control cells, we examined Golgi-localized protein synthesis and degradation rates in greater detail (**Figure 5A**). Notably, the data reveals a decreased synthesis of collagens COL6A1 and COL6A3 at 4 hours, preceding changes in collagen clearance rates. Notably, at prolonged ER stress, the increased clearance of collagen proteins including COL1A1, COL1A2, CO3A1, COL6A3, and COL12A1 coincides with the increased synthesis of Golgi protein maturation and collagen processing machineries including P4HB, TXNDC5, and GOLGA2, suggesting collagen clearance may be coupled to secretory pathway activity.

**Figure 5.**
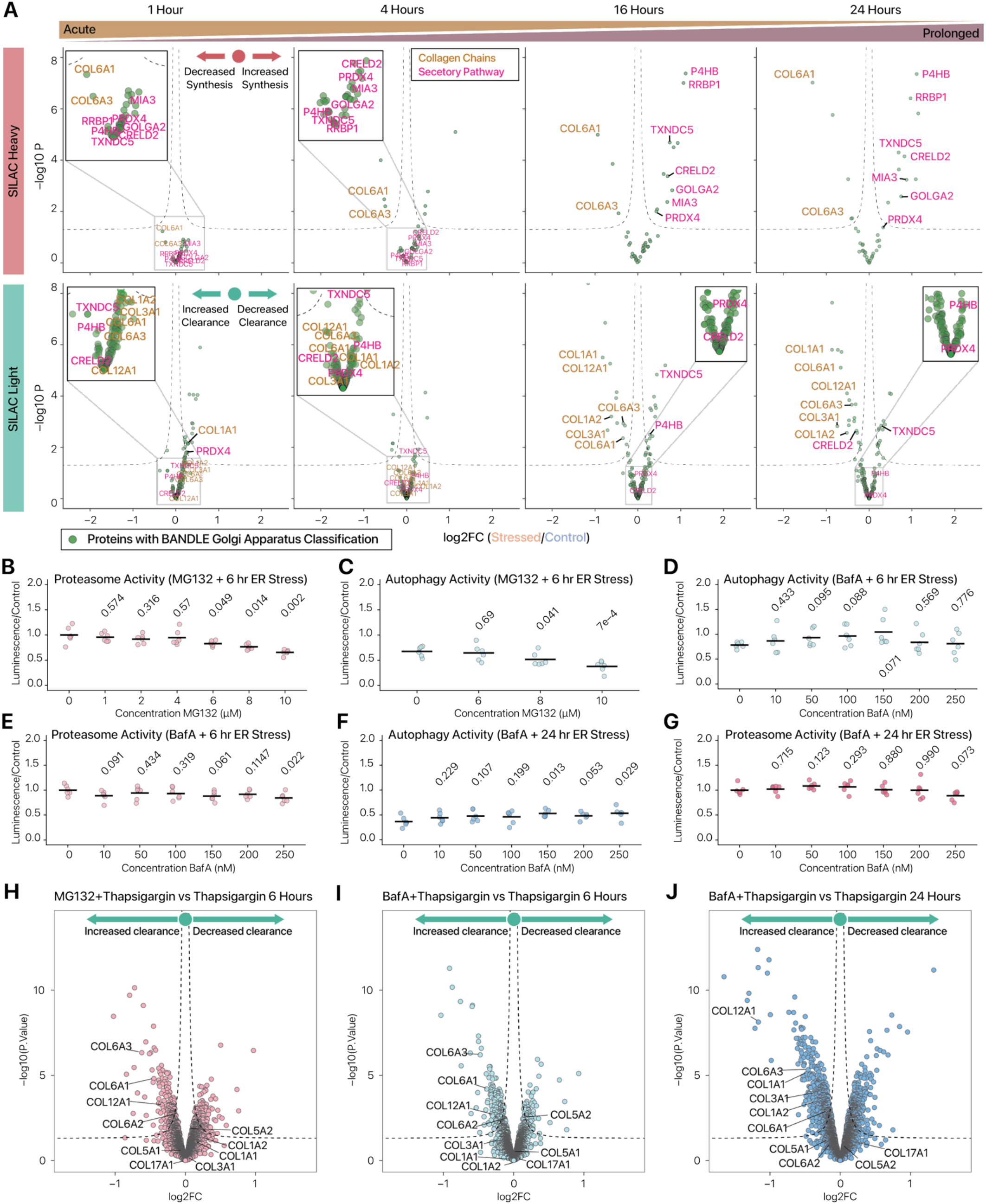
Golgi proteome remodeling suggest collagen initially aggregates and is subsequently cleared through secretion during ER stress. (A) Volcano plots generated by limma contrast fits for the heavy labeled and light labeled or unlabeled proteins predicted to reside in the Golgi apparatus at the 24 hour control for consistency. The y-axis represents the -log10 of the p-value and the x-axis represents the log fold change in intensity. The top set of volcano plots is for the heavy labeled proteins from the 1-hour timepoint (left) to the 24-hour timepoint (right). The bottom set of volcano plots is for the light labeled proteins from the 1-hour timepoint (left) to the 24-hour timepoint (right). Proteins labeled in pink are proteins associated with collagen processing. The proteins annotated in gold are collagen chains. (B) Assay of proteasomal degradative capacity under increasing concentrations of MG132 at 6 hours of 1 µM thapsigargin treatment. For panels B–G, all activity assay data points are plotted as a ratio to the average of the untreated condition. Significance was determined using a t-test comparing each concentration to the untreated condition. Each condition was performed with eight replicates. Solid line indicates the mean of the condition. (C) Assay of autophagic flux under increasing concentrations of Bafilomycin A at 6 hours of 1 µM thapsigargin treatment. (D) Assay of autophagic flux under increasing concentrations of Bafilomycin A at 24 hours of 1 µM thapsigargin treatment. (E) Assay of autophagic flux under increasing concentrations of MG132 at 6 hours of 1 µM thapsigargin treatment. (F) Assay of proteasomal degradative capacity under increasing concentrations of Bafilomycin A at 6 hours of 1 µM thapsigargin treatment. (G) Assay of proteasomal degradative capacity under increasing concentrations of Bafilomycin A at 24 hours of 1 µM thapsigargin treatment. (H) Volcano plots generated by limma contrast fits of SILAC light (unlabeled) proteins at 6 hours 1 µM thapsigargin exposure with 6 µM MG132 against 6 hours 1 µM thapsigargin exposure. The y-axis represents the -log10 of the p-value and the x-axis represents the log fold change in intensity. Arrows depict direction of increase vs decrease in clearance. (I) As in H, but for SILAC light (unlabeled) proteins at 6 hours 1 µM thapsigargin exposure with 150 nM BafA against 6 hours 1 µM thapsigargin exposure. (J) As in I but for SILAC light (unlabeled) proteins at 24 hours 1 µM thapsigargin exposure with 150 nM BafA against 24 hours 1 µM thapsigargin exposure.

To determine whether the collagens are degraded through transport to the lysosome-autophagy pathways, we examined significant differential localization of collagen chains from early to prolonged ER stress timepoints. The early timepoints (1 and 4 hours) are marked by the movement of multiple collagen chains from the Golgi towards a neighborhood adjacent to the “mitochondrial/chromatin” region of the spatial map (**Supplementary Figure S5**). As this area of the PCA maps to fractionation profiles of large complexes that sediment at low speed, this may represent aggregate formation or sequestration into specific membrane domains. Surprisingly, however, at the 16 hour timepoint the collagen chains instead move from the Golgi towards the plasma membrane rather than the lysosome, suggesting the collagen chains may be cleared via secretion rather than lysosome-autophagy degradation. To test this hypothesis, we inhibited proteasome and autophagy degradation at an intermediate ER stress timepoint and measured the clearance kinetics of the collagen chains.

Proteasome inhibition was achieved using 6 µM MG132, which in our hands significantly decreased proteasome activity without inducing autophagy (**Figure 5B–C**). At 6 hours, Bafilomycin A shows a dose-dependent effect on autophagy inhibition up to 150 nM without inhibition of proteasome activity (**Figure 5D–E**). The 150 nM dose remains effective at 24 hours of ER stress under a backdrop of proteolytic pathway remodeling (**Figure 5F–G**). We therefore used 6 µM MG132 and 150 nM Bafilomycin A to partially inhibit the ubiquitin-proteasome system and autophagy, respectively, while avoiding significant cross-reactivity on the other pathway. Unexpectedly, inhibition of autophagy did not abolish the increased collagen clearance at 6 hours of ER stress, arguing against autophagic removal (**Figure 5H**). Indeed, inhibition of either the proteasome or autophagy increased the clearance of collagen, suggesting there is an interaction between these degradation pathways and that collagen clearance is likely driven by increased secretion to the extracellular space or other unexpected mechanisms (**Figure 5I**). Examining inhibition of autophagy at a prolonged ER stress timepoint, 24 hours, we similarly observed increased collagen clearance suggesting this mechanism is conserved under prolonged ER stress (**Figure 5J**). Taken together, these data suggest that collagen clearance is associated with translocation across compartments, and that collagen processing and secretion may paradoxically increase under ER stress, suggesting prolonged ER stress in AC16 cells may trigger increased collagen processing through the secretory system as a proteostatic response. Moreover, the proteome-scale data highlight multiple overlapping degradation mechanisms including proteasome, lysosome-autophagy, and possible removal by secretion interact to orchestrate proteostasis as ER stress progresses.

### UFMylation-dependent ER-phagy acts in late ER stress cytoprotective protein clearance

We next examined the protein synthesis and clearance within the ER. The data revealed several ER-phagy-associated proteins to be up-regulated via both induced synthesis and decreased clearance at late ER stress timepoints (16-24 hours), including SEC24D, TMED8, TMEM214 and RPN1 (**Figure 6A**). This coincides with the degradation of multiple ER resident proteins, notably ERLIN1 and ERLIN2. As ERLIN1 and 2 are markers of specific ER lipid-raft domains and other domain markers (e.g., RTN3/4) are not prominently degraded, the data suggests ER-phagy may be activated under prolonged stress to degrade specific ER microdomains.

**Figure 6.**
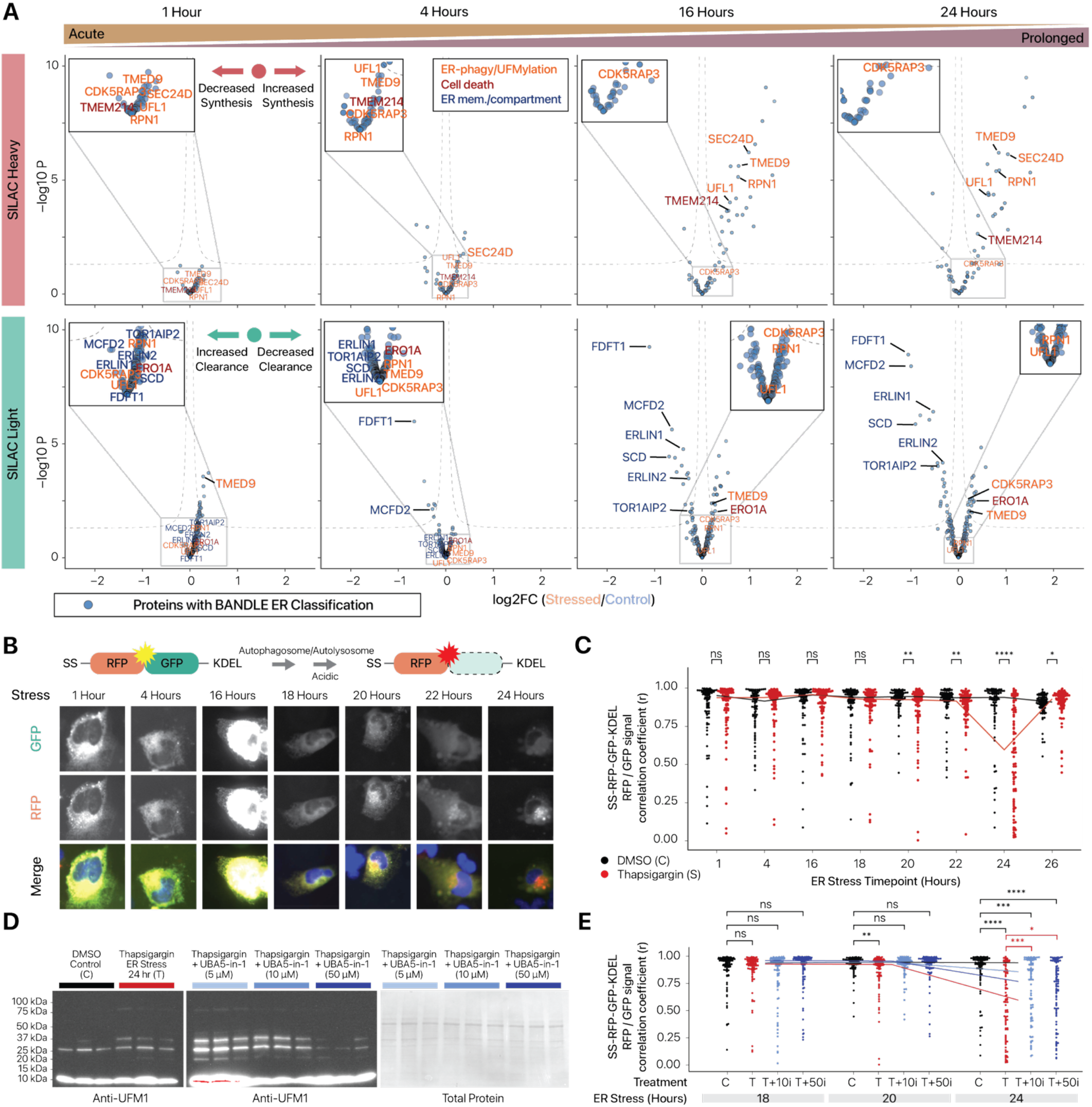
UFMylation induction coincides with increased ER lysosomal associated degradation during prolonged ER stress. (A) Volcano plots generated by limma contrast fits for the heavy labeled and light labeled or unlabeled proteins predicted to reside in the ER at the 24-hour control for consistency. The y-axis represents the -log10 of the p-value and the x-axis represents the log fold change in intensity. The top set of volcano plots is for the heavy labeled proteins from the 1-hour timepoint (left) to the 24-hour timepoint (right). The bottom set of volcano plots is for the light labeled proteins from the 1-hour timepoint (left) to the 24-hour timepoint (right). Proteins labeled in pink are proteins associated with collagen processing. Proteins labeled in orange are ER-phagy and UFMylation-related; proteins annotated in red are cell death related; proteins annotated in blue are ER membrane and microdomain proteins. (B) Top: Schematic of the ER-phagy assay. An ER localized construct containing a signal sequence, KDEL retention signal, GFP, and RFP emits a yellow light when localized at the ER. Upon ER-phagy activation, autophagosomes engulf portions of the ER and fuse with lysosomes. The acidic conditions of the lysosome denature the GFP but do not affect the RFP resulting in the emission of a red light. Bottom: High content imaging of a 1 µM thapsigargin time course on AC16 cells using ER-phagy assay. (C) Plots of the analysis of the high content imaging results for the correlation coefficient between the RFP and GFP signal (y-axis) over a time course of 1 µM thapsigargin treatment (x-axis). Each timepoint contains 90 cells screened for most average nuclear size of the cells imaged at that timepoint. DMSO treated samples (black) and thapsigargin treated samples (red) were compared at each timepoint using a Wilcoxon rank sum test after a Shapiro-Wilk test indicated the data deviated from normality. Solid line indicates the median of each condition. (D) Immunoblots for UFM1 for three replicates each of a 24-hour DMSO treated condition, 24-hour 1 µM thapsigargin treated condition, and 24-hour 1 µM thapsigargin treated conditions with indicated concentrations of UBA5-in-1 added 2 hours before experimental collection. UFM1 is at about 9 kDa and the upper bands are interpreted as UFM1 conjugated proteins. (E) High content imaging analysis of the correlation coefficient between the RFP and GFP signal (y-axis) over a time course of 1 µM thapsigargin treatment with indicated concentrations of UBA5-in-1. (x-axis). Each timepoint contains 90 cells screened for most average nuclear size of the cells imaged at that timepoint. Black brackets portray comparison to the DMSO treated condition. Red brackets portray comparison to the thapsigargin treated condition. UBA5-in-1 was added 2 hours before fixation. Each comparison was performed using a Wilcoxon rank sum test after a Shapiro-Wilk test indicated the data deviated from normality. Solid line indicates the median of each condition.

In our examination of ER proteome remodeling, both UFL1 and the macro-ER-phagy receptor CDK5RAP3 were differentially regulated at the late ER stress timepoints through increased synthesis and decreased degradation, respectively (**Figure 6A**). UFMylation is a post-translational modification that has been linked to ER-phagy (40–42), but the overall interactions between ER stress, UFMylation targets and ER-phagy activation are unclear. To verify ER-phagy induction in the stressed AC16 cells, we expressed an SS-RFP-GFP-KDEL reporter (43) followed by thapsigargin treatment (**Figure 6B**). The results confirm ER-phagy induction under prolonged ER stress, cumulating in peak activation at 24 hours (**Figure 6C**). The measured ER-phagy signal diminished after 24 hours, which may be due to RFP degradation or the synthesis of new dual-label reporters, although a down-regulation of ER-phagy perhaps concomitant with cell death cannot be ruled out. In parallel, UFM1 immunoblots show that ER stress leads to an accumulation of high-MW upper UFM1-conjugates, which are in turn suppressed by increasing concentrations of the UFMylation inhibitor UBA5-in-1 (**Figure 6D**). Moreover, prolonged-stress-induced ER-phagy as measured by the SS-RFP-GFP-KDEL reporters is significantly inhibited in the presence of the UFMylation inhibitor UBA5-in-1(**Figure 6E**), consistent with stress-induced increases of UFM1-conjugated proteins.

To assess whether ER-phagy activation is conserved in another cell type, we expressed the ER-phagy reporter in HeLa cells treated by thapsigargin. We observed a significant induction of ER-phagy at 22 and 24 hours in HeLa, albeit to a lesser degree than in AC16 cells (**Supplementary Figure S6A–B**). Imaging of stressed HeLa also revealed yellow (GFP+RFP) puncta that are not evident in AC16 cells, which is consistent with the presence of a population of autophagosome-engulfed ER portions not yet fused to lysosomes. Of note, cell viability assays show that these HeLa cells have lower viability than AC16 after 24 hours, suggesting a potential correlation between the degree of ER-phagy and the ability to mount viable stress response (**Supplementary Figure S6C**).

Multiple ER-phagy receptors have been found that facilitate autophagosome engulfment (44, 45), but the requirements of these receptors in different contexts is incompletely understood and debate remains on whether ER stress triggers ER-phagy or vice versa (40, 45). Prior work found that UFMylation of the ER resident protein CYB5R3 triggers CDK5RAP3-dependent ER-phagy (42). Hence, to evaluate that prolonged ER stress induced ER-phagy takes place via CYB5R3/CDK5RAP3, we expressed a CYB5R3-linked RFP-GFP ER-phagy reporter in AC16 cells. The CYB5R3 reporter reflected an increased ER-phagy upon ER stress; moreover, this induction is suppressed by the addition of UBA5-in-1, suggesting under prolonged-ER-stress-induced ER-phagy utilizes CYB5R3 UFMylation and engulfs CYB5R3-localized microdomains (**Figure 7A**). To assess whether other known macro-ER-phagy receptors are implicated in late ER stress, we expressed three ER-phagy receptor-linked reporters (TEX264-RFP-GFP, RFP-GFP-CCPG1, and RFP-GFP-FAM134C). Notably, ER-phagy activity linked to FAM134C was unchanged in stress but inhibited by the addition of UBA5-in-1 (**Figure 7B**), suggesting while UFMylation may promote ER-phagy via FAM134C, this receptor is unused in prolonged ER stress. Unexpectedly, UBA5-in-1 but not ER stress induced the TEX264 reporter signal (**Figure 7C**), whereas conversely, ER stress suppressed CCPG1 reporter activity, which is in turn rescued by UBA5-in-1 (**Figure 7D**). Taken together, these data indicate late ER stress involves different ER stress receptors, and that UFMylation has a complex and potentially bidirectional role that differentially affects ER-phagy receptors, where UFMylation inhibition suppressed the CYB5R3 and FAM134C reporters but activated TEX264 and CCPG1. It is not clear if UFMylation directly modifies ER-phagy receptors. To investigate this issue, we performed anti-UFM1 pulldown and mass spectrometry in control and 24-hour thapsigargin-treated AC16 cells. The analysis revealed a list of 334 proteins with significantly elevated pulldown in stressed cells (adj.P < 0.1 and abs(logFC) ≥ 0.5 vs. control; or exclusively identified in stressed cell pulldown), including UFM1 and UFL1 (**Supplementary Data S4**). Within this list of proteins, we did not identify ER-phagy receptors. However, 6 proteins (MARCKS, PCAS1, IRS2, MAFF, ATF6, IQGAP2) overlap with a recent study’s list of ER-phagy receptor interactomes in mouse PDAC cells (46), which may be the subject of future studies.

**Figure 7.**
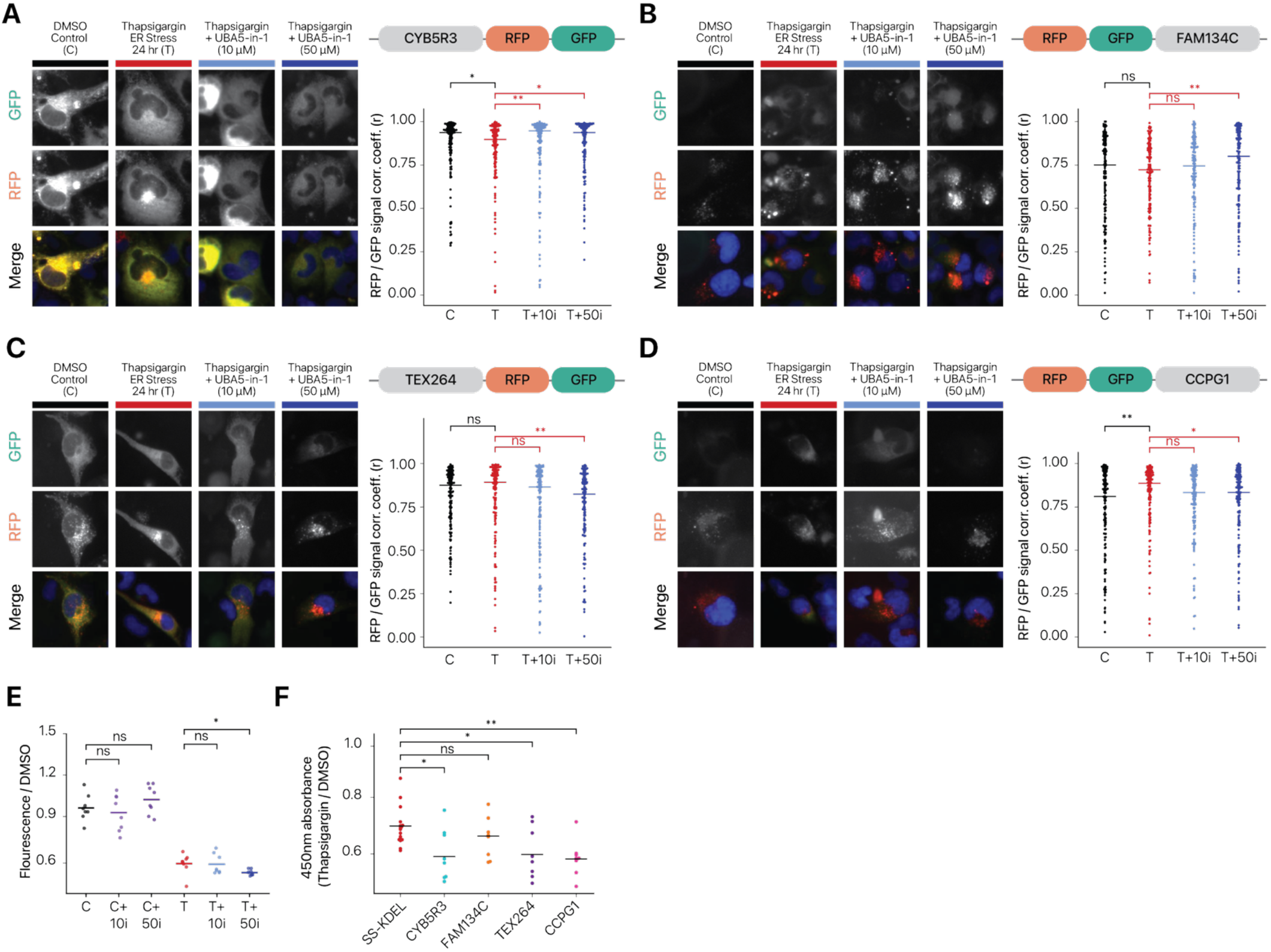
UFMylation selectively modulates ER-phagy receptor usage under prolonged ER stress. (A) Left: High content imaging of a CYB5R3-tagged RFP-GFP ER-phagy reporter at each condition performed in AC16 cells. Right: Plots of the analysis of the high content screening results for the correlation coefficient between the RFP and GFP signal (y-axis) over multiple conditions per construct (x-axis). Cells were exposed to DMSO or 1µM thapsigargin for 24 hours and UBA5-in-1 was added 2 hours before fixation. Each condition contains 150 cells screened for most average nuclear size of the cells imaged at that condition. Black brackets portray comparison to the DMSO treated condition. Red brackets portray comparison to the thapsigargin treated condition. Each comparison was performed using a Wilcoxon rank sum test after a Shapiro-Wilk test indicated the data deviated from normality. Solid line indicates the median of each condition. (B) As in E, but for the FAM134C-tagged RFP-GFP reporter. (C) As in E, but for the TEX264-tagged RFP-GFP reporter. (D) As in E, but for the CCPG1-tagged RFP-GFP reporter. (E) Plot of a resazurin viability assay with the fluorescence normalized to the DMSO control (y-axis) for 24-hour DMSO and thapsigargin treated conditions with and without indicated concentrations of UBA5-in-1 (x-axis). Solid line indicates the mean of eight replicates for each condition and each statistical comparison was performed using a t-test. (F) Plot of a CyQUANT XTT viability assay with the ratio of the absorbance at 450nm between the thapsigargin and DMSO condition of each construct (y-axis) for each construct (x-axis) in AC16 cells after 24 hours of treatment. Statistics were performed using a t-test between each construct to the KDEL. The KDEL readings were performed with 16 replicates, and every other construct was performed with eight replicates. Solid line indicates the mean of the construct and each statistical comparison was performed using a t-test.

Finally, to evaluate whether UFMylation causally affects stress remediation, we examined the effect of UBA5-in-1 on cell survival. UBA5-in-1 synergistically increased cell death in ER stressed cells but not normal cells, suggesting ER-phagy in prolonged ER stress is cytoprotective (**Figure 7E**). These results align with previous results reporting decreased cell death with the overexpression of UFMylation machinery (41). Because the expression of the ER-phagy receptor-linked reporters themselves may stimulate ER-phagy, we also examined whether their expression impacted cell viability relative to the KDEL reporter. Unexpectedly, comparing the ratio of viability between the stressed and unstressed cells showed a modest decrease in viability for reporter expressing cells under stress (**Figure 7F**), suggesting the reporters are not by themselves cytoprotective. This prompts a model where the cytoprotective properties of UFMylation arise from its action on specific ER-phagy receptors, including by inhibiting TEX264 and CCPG1 activity, which may be explored in future work.

## DISCUSSION

This work describes SPLAT-TR, a mass spectrometry-based method to attain global maps of protein synthesis and degradation across cellular compartments. SPLAT-TR uses a dimension reduction informed approach to simplify the number of fractions in ultracentrifugation preparation while retaining spatial resolution, allowing spatial information to be encoded alongside a TMT-SILAC design that disaggregates synthesis and clearance estimates. Applying SPLAT-TR to early and prolonged ER stress models, we reveal previous underappreciated aspects of ER stress response, including widespread protein translocation, protein-specific synthesis and clearance changes, and the differential utilization of degradation pathways targeting specific proteins across organelles.

UPR signaling branches show differential attenuation kinetics (5, 10, 47), as well as considerable target overlaps and mutual regulation (9, 48, 49), together producing a complex network of downstream proteostatic responses. Our understanding of the molecular consequences of UPR is continuing to emerge; for instance, proteomics profiling has discovered an expanded regulon of ∼200 ER stress response genes involving a rewiring of metabolic phenotypes (11); whereas another recent work suggests UPR induces the synthesis of splice factors to activate a conserved splicing network (50). In prior work, we found multiple cell surface channels and receptors selectively translocating to and from the cell membrane under ER stress (18). Here we significantly expanded our investigation to incorporate protein synthesis and clearance changes across early and prolonged ER stress timepoints. Spatiotemporal processes in particular underlie the early proteomic remodeling in 1–4 hours of ER stress, which were poorly characterized from protein abundance measurements alone (11).

Protein homeostasis requires the clearance of existing proteins through multiple protein degradation pathways. Cellular compartments feature specialized degradation machineries that adapt to local proteostatic needs, but details on which proteins are targeted by which pathways are still emerging. Within the secretory pathway, ERAD serves to remove newly synthesized misfolded proteins (51, 52) whereas other proteins, complexes, and aggregates may be removed by autophagy (53–56). Defining the respective contributions and clients of these pathways has been complicated by their interactions, where they may target overlapping proteins (57), ERAD machinery may be targeted by autophagy, and ERAD dysfunction may induce autophagy (58, 59).

Investigating synthesis and degradation changes across stress response timepoints, we find several lines of evidence suggesting a dynamic dependency on differing protein clearance pathways over the course of ER stress. Differential localization of proteins toward lysosomes is predominant only in late timepoints, whereas total proteome capacity remained relatively level in early and mid ER stress but collapsed during late ER stress time points. Prior copy number and kinetics calculations suggest there is limited excess in proteasome capacity even in cells under basal condition (60). The drop in capacity coupled to increased degradation demand points strongly to a shift in late ER stress from initial proteasome degradation toward relying on other degradation pathways as the main strategy to maintain proteostasis and viability. The decrease in proteasome capacity coincides with the increase in lysosomal degradation of secretory proteins and non-proteasomal clearance of collagen. In particular, our spatial and proteomic remodeling data suggest collagen secretion may be an alternative mechanism of collagen removal in prolonged stress, which is supported by the lack of clearance inhibition during inhibition of autophagy or the proteasome. In neurodegenerative disease models, mutant HTT is actively secreted under stress to reduce aggregate formation (61). Our results here suggest secretion may present a general stress response mechanism that applies to multiple Golgi-localized collagen chains. While prior work has shown that collagen clearance occurs via autophagy, the discrepancy may be due to differences in cell line or stressor; e.g., prior work suppressing of the collagen folding chaperone HSP47 may induce misfolding and alternative clearance mechanisms (39, 62).

Also coinciding with the decrease in proteasome capacity is an induction in the synthesis of ER-phagy-related proteins, suggesting they play a role in prolonged ER stress response. In the literature, ER-phagy has been described to both be cytoprotective for stressed cells (41, 63, 64) while excessive ER-phagy may promote cell death (65, 66). These prior investigations of cell viability have largely focused on the contributions of FAM134B. To our knowledge, our characterization of the effect on cell viability for the other ER receptors has not been reported and suggests receptors may have opposite roles in cell death and survival and may further have distinct regulation modalities through UFMylation. The functional diversification and significance of ER-phagy receptors remains a subject of intense research, but it is known that they can drive the engulfment of different cargos (67, 68). In parallel, we show that ER-phagy in AC16 cells is largely dependent on UFMylation, whereas inhibition of UFMylation decreases cell viability in our hands, agreeing with prior work showing overexpression of UFMylation machinery is cytoprotective (41). Future work may further investigate the distinction between ER-phagy driven by UFMylation and receptor expression, as well as the effect on cell viability.

The ER is the largest organelle in the cell and forms extensive contacts with other cellular compartments (69). Whereas early work has focused on the isolated effect of ER stress on proteostasis of ER proteins, our proteome-wide investigation joins a chorus of recent work that highlights the cell-wide coordination of proteostasis across organelles. The propagation of proteostatic signals likely involves in part the re-localization of multiple proteins, including those related to endocytosis and vesicular transport (4 hours) as well as nuclear actin and chaperone networks (16–24 hours). As the ER plays important roles in endosome maturation (70–72), the early translocation changes nominate endocytic suppression as a potential aspect of early ER stress. In parallel, we also observed the re-localization of multiple subunits for the ARP2/3 complex, an actin polymerization regulator, toward the nucleus. Nuclear ARP2/3 plays important roles in DNA damage repair (30) and nuclear actin has been associated with transcriptional regulation and chromatin restructuring (73, 74). To our knowledge, a stress-induced ARP2/3 re-localization to the nucleus has not been described, supporting the utility of SPLAT-TR to nominate new proteomic remodeling events. Future work may explore the mechanism through which the re-localization is regulated and the effect it has on actin organization.

The close contact between the ER and mitochondria has particularly been implicated in ER stress response (75). UPR signaling proteins including PERK and IRE1a have been found to be enriched at MAM (76, 77). MERCS have been observed to increase during early ER stress to facilitate calcium transfer and ATP production (35), but are also implicated in mitochondrial calcium overload and cell death (76–78). Here, we show that this biphasic response from adaptive to maladaptive ER stress is underpinned by time-dependent changes in protein synthesis and clearance programs, where an initial increase in the synthesis of bioenergetics proteins at 1 hour quickly subsides and becomes replaced by the coordinated accumulation of mitochondrial protein quality control machineries. The results suggest a model where increased ATP production and ROS production in early timepoints may serve to propagate initial ER stress to mitochondrial proteostatic challenge and exacerbate overall cellular proteostatic load.

Finally, the SPLAT-TR design also provides insights into spatial resolution achievable through protein correlation profiling/localization of organelle proteins (PCP/LOP) methods. It is generally held that greater numbers of separation steps must afford greater granularity in spatial resolution. A recent work examined the trade-off in ultracentrifugation fraction counts to balance spatial resolution with data incompleteness but found that removal of some fractions led to lower resolution, although it is unclear how the fractions were selected for removal (79). In contrary, by grouping the fractions in a cell type based on similarity, we showed that we maintained or even enhanced resolution in the compressed fractionation approach. This result may point to a limit in the separation power of organelle fractionation when based on the same physiochemical principles, and that future work may compare enhancing separation using additional physiochemical properties.

### Limitations of the study

SPLAT-TR requires TMT and SILAC dual labeling which adds to the cost of experiment. The number of available TMT channels also places a limit to the number of spatial fraction and temporal timepoints that can be compared in one experiment under the TMT-SILAC design, which may be circumvented by ongoing advances that explore the use of label-free quantification (LFQ) data from data-independent acquisition (DIA) mass spectrometry to achieve comparable spatial resolution while foregoing TMT labeling (79, 80); however, such approaches currently remain less common in subcellular proteomics and are an area of ongoing development. Secondly, while the rationale behind TMT-SILAC quantification of protein synthesis and clearance is well established (24, 25), we did not attempt to independently verify protein synthesis such as using BONCAT or puromycin-based methods (50). The contribution of cell proliferation to protein clearance was also not investigated. Lastly, we are unable to attribute the full ER-phagy signal to any one receptor. CYB5R3 was the only ER-phagy-linked protein whose engulfment is induced by ER stress in AC16 cells in our hand, but our study is limited by the effect of overexpression on non-stressed ER-phagy activity and thus other receptors likely also participate in ER stress induced ER-phagy. ER-phagy can also occur via vesicle budding off the ER and direct delivery to lysosomes as opposed to autophagosome engulfment of ER segments, which may explain some reporter results.

## Resource Availability

### Lead Contact

Requests for further information and resources should be directed to and will be fulfilled by the lead contact, Edward Lau.

### Materials Availability

This study did not generate new unique reagents.

### Data and Code Availability

Raw mass spectrometry data for the spatiotemporal proteomics data sets are available on ProteomeXchange under Accession for PXD082623 with access token rkdY2gEVy88G.

Raw mass spectrometry data for UFM1 pulldown experiments are available on ProteomeXchange under Accession PXD083294 with access token 7fXfljoLAdCu

The R code for data analysis and figure generation is available on GitHub under https://github.com/LorenaAlamillo/SPLATTR-Publication-Figure-Generation. Riana is available on GitHub under https://github.com/ed-lau/riana. pyTMT is available on GitHub under https://github.com/Lau-Lab/pytmt.

## Acknowledgments

This work was supported in part by NIH National Institute of General Medical Sciences (NIGMS) Diversity Supplement Program support to L.A.; NIGMS award R35GM146815 and National Heart, Lung, and Blood Institute (NHLBI) grant R00HL144829 to E.L.; and NHLBI grant R01HL141278, NIGMS grant R01GM144456, and the University of Colorado School of Medicine Advanced Proteomics Infrastructure (API) funds to M.P.Y.L. T.A.M. was supported by National Institutes of Health grants R01HL171711 and HL181226 and an American Heart Association Collaborative Sciences Award (24CSA1255857). J.G.C. received support from National Institutes of Health project grant T32GM158468. Imaging data were acquired in part at the Advanced Light Microscopy Core facility of the NeuroTechnology Center at the University of Colorado Anschutz Medical Campus, which is supported in part by the Diabetes Research Center Grant (NIH P30DK116073). Usage of the Zeiss LSM 980 was supported by NIH S10OD036220.

## Author Contributions

Conceptualization, L.A., M.P.Y.L. and E.L.; methodology, L.A., T.P., M.P.Y.L. and E.L.; investigation, L.A., T.P., A.B., M.C.J., E.A., J.C, D.C.M.N.; writing – original draft, L.A. and E.L.; writing – review & editing, L.A., M.P.Y.L. and E.L.; funding acquisition, T.A.M., M.P.Y.L., and E.L.; resources, T.A.M., M.P.Y.L., and E.L.; supervision, M.P.Y.L. and E.L.

## METHODS

### AC16 cell culture, heavy isotope labeling, ER stress induction

AC16 cells (Millipore) at passage 8 were cultured in DMEM/F12 supplemented with 10% FBS and maintained at 37°C with 5% CO_2_ and 10% O_2_

. To perform the isotopic labeling, the culture media was replaced with SILAC DMEM/F12 (Thermo Scientific) deficient in both L-lysine and L-arginine supplemented with 1% dialyzed FBS and heavy amino acids at concentrations 0.499 mM for ^13^C_6_^15^N_2_ L-Lysine-2HCl and 0.699 mM for ^13^C_6_^15^N_4_ L-Arginine-HCl (Thermo Scientific). To initiate ER stress, 1 µM thapsigargin (SelleckChem) was added concurrently to the initiation of isotopic labeling for each timepoint of 1 hour, 4 hours, 16 hours, and 24 hours alongside a non-treated control.

### Cell harvesting, differential centrifugation, and isobaric labeling

The AC16 cells were harvested with trypsinization, washed 3× with room temperature PBS, and resuspended in a detergent free gentle lysis buffer (0.25 M sucrose, 10 mM HEPES pH 7.5, 2 mM magnesium acetate). 3 mL of the resuspension at a time was lysed using an Isobiotec ball-bearing homogenizer with a 16 µM clearance size using ∼30 passages through the chamber. Cell rupture was verified at ∼80% with trypan blue. The resulting lysate for each experimental collection was centrifugated 3 times at 4°C in a swinging bucket centrifuge at 200 × g for 5 min, removing the pelleted unlysed cells and continuing with the supernatant each time. The final supernatant was ultra centrifugated according to Table 1 to generate the 4 ultracentrifugation pellets and supernatant 5.

The supernatant generated in the final spin was separated and all pellets were stored at –80°C until proceeding. The proteins in the supernatant were precipitated in 4× the volume of cold acetone overnight at –20°C. The precipitate was pelleted by centrifuging at 13,000× g for 10 min at 4°C generating pellet 5. The acetone was removed and the pellet was allowed to dry for ∼30 min before resuspension in a resolubilization buffer consisting of 8 M urea, 50 mM HEPES pH 8.5, and 0.5% SDS with 1x Halt Protease and Phosphatase Inhibitor Cocktail (Thermo Scientific). Pellets from the ultracentrifugation fractions 1 to 4 were resuspended in RIPA buffer with 1x Halt Protease and Phosphatase Inhibitor Cocktail (Thermo Scientific). Resuspended pellets 1 through 5 were sonicated in a Bioruptor sonicator with settings 20× 30 s on 30 s off at 4°C. The sonicated samples were then centrifugated at 14,000 ×g for 5 min to remove insoluble debris. The protein concentration of the supernatant was measured with Rapid Gold BCA (Thermo Scientific).

Digestion and isobaric labeling follows that of the published iFASP TMT protocol (81). For each replicate, the 1-hour and 16-hour timepoints were digested in parallel as were the 4-hour and 24-hour timepoints. Pierce Protein Concentrators PES, 10 K MWCO were prewashed with 100 mM TEAB and 250 µL of 8 M urea was loaded onto the filter. A total of 100 µg protein per fractions 1 to 3 were individually added to their own concentrator. Fractions 4 and 5 were combined at a mass ratio of 2:1 fraction 4 to 5 for a total of 100 µg and loaded on to a concentrator together. The samples were washed with 250 µL 8 M urea to denature proteins and remove SDS and twice with 300 µL 100 mM TEAB to remove the urea. Washing was followed with reduction and alkylation using a 30 min incubation at 37°C with TCEP and CAA in the dark. The TCEP and CAA were removed through centrifugation and 3 washes with 300 µL 100 mM TEAB. After the final wash, the samples were digested with mass spectrometry grade trypsin (Promega) overnight at 37°C at a ratio of 1:50 enzyme:protein. TMTpro-16plex isobaric labels (Thermo Scientific) were equilibrated to room temperature and reconstituted in 20 µL LC-MS grade anhydrous acetonitrile. For each replicate, the 1 hour and 16 hours timepoints were multiplexed together as were the 4 hours and 24 hours timepoints. In each round of labeling, labels were randomly assigned to each fraction (**Supplementary Table S2**) with a random number generator in Excel to mitigate possible batch effects. The resuspended isobaric labels were added to the samples and incubated at room temperature for 1 hour with shaking. 1 µL 5% hydroxylamine was added to each sample and incubated at room temperature for 30 min with shaking to quench the reaction. Peptides were eluted in two rounds with an initial centrifugation step followed by the addition of 40 µL 50 mM TEAB to the filters to elute with an additional round of centrifugation. All 16 labeled fractions per round of labeling were combined and mixed well before separating 100 µg calculated by input mass over the final volume. Aliquots were dried with speed-vac and stored at –80°C.

### Liquid chromatography and tandem mass spectrometry

RP-LC was performed on each 100-µg multiplexed sample to obtain additional sample depth. Each 100-µg aliquot was reconstituted in 50 µL of 20 mM ammonium formate pH 10 in LC-MS grade water (solvent A) and injected into a Jupiter 4 -µm Proteo 90-Å LC Column of 150 × 1 mm on an Ultimate 3000 HPLC system. The gradient was performed with a flow rate of 0.1 mL/min as follows:, 0–30 min: 0%–40% Solvent B (20 mM ammonium formate pH 10 in 80% LC-MS grade acetonitrile); 30–40 min: 40%-80% Solvent B; 40–50 min: 80% Solvent B. Fraction collection occurred every minute and samples with little intensity were pooled resulting in a total of 16 peptide fractions. The fractions were dried on the speed-vac and stored until further processing.

Fractions were reconstituted in 10 µL each of pH 2 MS solvent A (0.1% formic acid) and analyzed with LC-MS/MS on a Thermo Q-Exactive HF orbitrap mass spectrometer coupled to an LC with electrospray ionization source. Peptides were separated with a PepMap RSLC C18 column 75 µm x 15 cm, 3 µm particle size (Thermo Scientific) using a 90-minute gradient from 0 to 100% pH 2 MS solvent B (0.1% formic acid in 80% LC-MS grade acetonitrile). Full MS scans were acquired with a 60,000 resolution. A stepped collision energy of 27, 30, and 32 was used, and MS2 scans were acquired with a 60,000 resolution and an isolation window of 0.7 m/z.

### Mass spectrometry data processing

Mass spectrometry raw data were converted to mzML format using ThermoRawFileParser v.1.7.3 (82) then searched against the UniProt Swiss-Prot human canonical and isoform protein sequence database (retrieved 2024-04-26) using Comet v.2023_01_rev0 (83). The fasta database was further appended with contaminant proteins using Philosopher v4.4.0 (total 42,402 forward entries). The search settings were as follows: peptide mass tolerance: 10 ppm; isotope error: 0/1/2/3; number of enzyme termini: 1; allowed missed cleavages: 2; fragment bin tolerance: 0.02; fragment bin offset: 0; variable modifications: TMTpro-16plex tag +304.2071 for TMT experiments, and lysine +8.0142, arginine +10.0083 for all SILAC experiments; fixed modifications: cysteine + 57.0214. The search results were further reranked and filtered using Percolator v3.05 (84) at a 1% FDR.

### TMT labeling assignment, isotope correction, and purity assessment

TMT label quantification was performed using pyTMT (85) on the Comet/Percolator output. pyTMT extracts peptide intensities from the MS2 encoded information and corrects for cross contamination in the TMT tags using the batch contamination data sheet (**Supplementary Table S3; Supplementary Table S4**). TMTpro-16 plex lots were #XL348283 for replicate 1 and 2 and #YB367250. for replicate 3. The true channel matrix is calculated from the observed channel intensity and impurity matrix using the non-negative least square algorithm in scipy (86).

To determine the protein distribution profiles over the set of fractions, peptide levels intensities were summed to the protein level. To obtain a representative signal, peptides mapping to more than one non-isoform UniProt protein accession were discarded. Protein isoforms mapping to the same UniProt protein accession were discarded. Peptides mapping to a canonical and non-canonical isoform without a unique peptide were assigned to the canonical. Only isoforms with unique mapping peptides were included in this study. Light and heavy peptides were individually summed with the by labeling peptides/proteins containing a heavy SILAC modification with ‘_H’ in the data tables and with “*” in figures. The proteins are then column and row normalized before further analysis.

### Heavy and light peptide quantification

Heavy and light peptide quantification was performed by measure the area under the curve using Riana v0.8.0 (87). Riana integrates the peak intensity encompassing the first and last MS2 scan where the peptide is confidently identified for the chosen isotopic peaks. For this experiment, Riana was set to recognize a 25-ppm error of the light (+0) and heavy (+8, +10) peptide peaks over a 20 -second retention time window. Because the sample information was encoded in the MS2 level information, we used the MS2 level information to convert the total area under the curve for the peptides to the intensity of the peptide per sample. We used the pyTMT output to calculate the ratio contribution of each sample to the total MS2 peptide intensity. The Riana output peptides were mapped to protein groups as they were for the pyTMT output allowing for peptides to be mapped between the different outputs. The AUC was converted to MS1 intensities per heavy or light version of the peptides with heavy labeled peptides annotated with a ‘_H’ in the data tables and with “*” in figures.

### Comparing synthesis and clearance

The MS1 intensities per heavy or light peptides were summed to the protein level and normalized with variance stabilization normalization before analysis with limma (88). The data was corrected for batch and replicate to compare the stressed condition to the control. This was separately performed for the heavy protein, light protein, and the sum of the heavy and light protein. Subcellular analysis of heavy and light proteins was performed by isolating the proteins assigned to the subcellular compartment of interest. Assignments at the 24-hour control timepoint were used for all timepoints to keep the proteins compared consistent.

### Subcellular localization classification

Subcellular localization predictions were performed between all replicates of the control and all replicates of the thapsigargin treated at each timepoint using the pRoloc (89) and the BANDLE (26) packages. To train the supervised learning model for spatial classification, we began with subcellular markers derived from a prior data set generated from human U-2 OS osteosarcoma cells by the Lilley lab (20). These initial markers were filtered for the proteins intersecting with the AC16 dataset and further curated to account for cell type-specific marker expression. Proteins were pruned from the marker list if they showed deviation from the average distribution in any of the timepoints, conditions, or replicates. Furthermore, proteins were added to the marker list if they remained consistently within a 95% confidence interval Mahalanobis distance ellipse in every timepoint, condition, and replicate PCA. The added markers were cross referenced to Gene Ontology (GO) Cellular Compartments to ensure there were no conflicting annotations. The cytoskeleton markers showed poor agreement across replicates, so we removed the cytoskeleton from the subcellular compartments analyzed. We also removed chromatin from the subcellular compartments analyzed as the distribution profiles for the chromatin markers largely overlapped with those of the mitochondrion markers leading to multiple reports of false translocations.

Next, the subcellular markers were visually assessed and manually pruned for the markers with a protein distribution profile differing from that of the majority of the markers for each subcellular compartment. New markers were assigned by whether they were consistently within a Mahalanobis distance ellipses with a chi-squared of 0.99 at every condition/timepoint/replicate. Newly added markers were cross referenced with UniProt annotations for subcellular location and cellular function; proteins with conflicting annotations were removed. For differential localization analyses we used the Markov-chain Monte-Carlo (MCMC) and non-parametric model in BANDLE to find unknown protein classification and evaluate differential localization probability. MCMC parameters are 9 chains, 10,000 iterations, and 5000 burn-in, 20 thinning, seed 42; convergence of the Markov chains is assessed visually by rank plots (90). QSep was performed using the QSep function with the Bandle output in msn format. QSep reports separation values for each compartment to every compartment in the dataset. These separation values were averaged for each compartment within replicates and standard deviations and means between replicates were visualized.

### Fluorescent imaging validation

N-terminally snap-tagged constructs were designed into a plasmid cloning deoxyribonucleic acid (pcDNA) 3.1(+) backbone for each protein. AC16 cells were plated on a 6-well plate and incubated with conditions described above. Two wells per construct were transfected using lipofectamine 3000 (Thermo Scientific) with 2.5 µg of construct per well according to kit instructions. Opti-mem (Thermo Scientific) was used to combine the components of the reactions. The transfection media was removed following 4 hours of incubation and replaced with culturing media. The next day the transfected cells were passaged using trypsinization and pooled for each construct. The cells were plated on a 96-well plate at a density of 8,000 cells for the early timepoints and 4,000 cells for the later timepoints with early and late timepoints on separate plates. Immediately before starting treatments, the cells were labeled with soluble N-ethylmaleimide-sensitive factor attachment protein (SNAP)-Cell Oregon Green (New England Biolabs) at a concentration of 4 µM for 30 min at 37°C subsequently the media was removed and the cells were washed with media 3 times for 30 min incubations at 37°C. Eight wells per construct were treated with either a DMSO vehicle control or thapsigargin and incubated for 1 hour, 4 hours, 16 hours, or 24 hours. At the end of the incubation period, the cells were fixed with 4% PFA in PBS (Thermo Scientific) for 15 min at room temperature, washed 3 times with PBS, permeabilized with 0.01% triton in PBS for 15 min at room temperature, and once again washed 3 times with PBS. The cells were labeled with deep red high content screening cell mask (Thermo Scientific) at a ratio of 1:5000 in PBS for 30 min at room temperature protected from light and subsequently washed 3 times with PBS. Cells were then labeled with NucBlue Fixed Cell Stain ReadyProbes reagent (Thermo Scientific) diluted into 1mL PBS per 2 drops reagent for 5 min room temperature protected from light. The cells were washed 1 time with PBS before the addition of 0.1% sodium azide in PBS in each well. The cells were stored at 4°C until further processing.

Imaging was performed using the Cell Insight CX7 LZR Pro High Content Screening Platform (Thermo Scientific) using the preprogrammed single translocation assay which allows for the creation of two overlayed masks, an inner circle and outer ring. For the nuclear translocation assay, the nucleus was defined by the inner circle with the outer ring designated as the cytoplasm. The high-content analysis platform reports the ratio of the fluorescence intensity within the inner circle to the intensity within the outer ring. These results were further processed in R with 50 cells selected per condition by the most average nuclear size. The results were plotted and statistics were performed using Wilcoxon rank sum test comparing the control and thapsigargin treated at each timepoint as a Shapiro-Wilk test indicated the data deviated from normality. For the membrane translocation assay, the whole cell stain was used to assign the inner circle as the cytoplasm and the outer ring as the free edge of the plasma membrane. The results were further processed in R to calculate the intensity ratio of the membrane (ring) to the cytoplasm (circle) and filter for the top 50 cells expressing the protein of interest for each timepoint. The results were plotted and statistics were performed using Wilcoxon rank sum test comparing the control and thapsigargin treated at each timepoint as a Shapiro-Wilk test indicated the data deviated from normality.

### Seahorse assay

AC16 cells (Millipore) at passage 6 were passaged onto a XF96 Cell Culture Microplate (Agilent) at a density of 10,000 cells per well, excluding wells along plate edges, and let incubate overnight. Five wells per timepoint/condition were treated with either 0.1% DMSO vehicle control or 1 µM thapsigargin in normal media for the indicated timepoints. Seahorse XF DMEM (Agilent) was supplemented with 17.5 mM glucose (Agilent), 2.5 mM glutamine (Agilent), and 0.5 mM pyruvate (Agilent) to create assay medium. A Seahorse XFe96/XF Pro Sensor Cartridge (Agilent) sensor cartridge was hydrated overnight in Seahorse XF Calibrant (Agilent) and loaded with Seahorse XF Cell Mito Stress Test Kit (Agilent) constant-compound-variable-loading solutions resuspended in Seahorse XF DMEM assay medium (Agilent) according to manufacturer instructions. At the end of the treatment period, cells were washed with Seahorse XF DMEM assay medium (Agilent) three times, leaving 20 µL in the well after each wash, and incubated in a total of 180 µL of XF medium for 45 mins at 37°C in a non-CO2 incubator. Data was collected using a calibrated Seahorse XF96 Analyzer over a period of approximately 75 minutes, during which oligomycin, FCCP, and antimycin A + rotenone were sequentially injected at final concentrations of 1.5 uM, 1 uM, and 0.9 uM, respectively. The data was normalized by total protein content of each well measured using a Pierce BCA protein assay kit (Thermo Scientific). Data quality was controlled using Wave Controller 2.6, and analysis was performed using the Seahorse Analytics software (Agilent). Mitochondrial activity was interpreted from each respective compound treatment and averaged over technical replicates. Statistical significance between conditions and timepoints was determined using ANOVA followed by pairwise comparisons with Holm adjustment. Bar plots were generated comparing each thapsigargin treatment timepoint with the pooled DMSO controls across all timepoints.

### Fluorescence resonance energy transfer assay

AC16 cells (Millipore) at passage 6 were plated onto a 6-well plate and cultured as described above. At about 80% confluency cells were transfected with 1.25 µg of SEC61B-CFP and 1.25 µg of TOM20-YFP each within a pcDNA3.1(+) backbone. The next day the cells were passaged onto glass 8-well slides (ibidi) at a density of 25,000 cells per well. Four wells per timepoint/condition were treated with either DMSO vehicle control or thapsigargin for the indicated timepoints. At the end of the incubation period, the cells were fixed with 4% PFA in PBS (Thermo Scientific) for 15 min at room temperature, washed 3 times with PBS for 5 mins each wash, and mounted using ProLong Glass Antifade Mountant (Invitrogen). Ten cells per well were imaged using a Zeiss LSM 980 using the preprogrammed excitation and emission profiles for tagCFP and tagYFP. CellProfiler v4.2.8 was used to identify CFP and YFP overlapping regions within each cell and normalized FRET was calculated using the CFP, YFP, and FRET integrated intensities. FRET was normalized for expression of YFP and CFP using Equation 10 in (91) and adjusted for bleedthrough of the donor and acceptor channels using equation 11 in (92). The amount of positive FRET regions per cell was calculated and compared between conditions at each timepoint using a Wilcoxon rank sum test.

### Proteasome degradative capacity assay

AC16 cells (Millipore) at passage 9 plated into a 96-well plate at a density of 5,000 cells per well and were cultured as described above. The next day, eight wells per timepoint were exposed to 1 µM thapsigargin before addition of the Proteasome-Glo (Promega) reagent as specified by manufacturer. Luminescence was detected using the Spectral Max iD5 (Molecular Devices) luminescent setting. For inhibitor concentration experiments, Bafilomycin A1 (Sigma Aldrich) or MG132 (Fisher Scientific) was added at indicated concentrations 2 hours before collection.

### Autophagic flux assay

AC16 cells (Millipore) at passage 9 were plated into a 96-well plate at a density of 7,000 cells per well and were cultured as described above. The next day, the cells were transfected with 0.025 µg of the microtubule-associated protein 1A/1B-light chain 3 (LC3) HiBit Reporter from the Autophagy LC3 HiBit Reporter Assay System (Promega) using lipofectamine 3000 (Thermo Scientific). After overnight incubation, eight wells per timepoint were exposed to 1 µM thapsigargin before addition of the Nano-Glo HiBit Lytic Reagent (Promega) as specified by manufacturer. Luminescence was detected using the Spectral Max iD5 (Molecular Devices) luminescent setting. For inhibitor concentration experiments, Bafilomycin A1 (Sigma Aldrich) or MG132 (Fisher Scientific) was added at indicated concentrations 2 hours before collection.

### Sample processing and LC-MS/MS for protein degradation inhibitor experiments

AC16 cells (Millipore) at passage eight were plated into 6-well plates and, once the cells were approximately 80% confluent, cultured for isotopic labeling as described above. 1 µM thapsigargin (SelleckChem) was added concurrently to the initiation of isotopic labeling for 6 or 24 hours. For inhibited conditions, MG132 (Fisher Scientific) was added at 6 µM or Bafilomycin A1 (Sigma Aldrich) was added at 150nM 2 hours before collection. Cells were harvested with trypsin, pelleted at 300 ×g for 3 min and washed with 1 mL PBS by inverting 2×. Pellets were snap frozen using liquid nitrogen and stored at −80C until further processing. As described above, cells were resuspended, sonicated, digested, and isobaric tag labeled (**Supplementary Table S5**). Samples were further processed using RPLC and analyzed with LC-MS/MS on a Thermo Orbitrap Exploris 480 mass spectrometer coupled to an LC with electrospray ionization source using previously described settings. Raw files were converted following pipeline outlined above through heavy and light peptide quantification and comparing synthesis and clearance. TMT contaminant matrix correction for lot #YB367250 was used.

### ER-phagy reporter assay

The ER-phagy reporter construct SS-mRFP-eGFP-KDEL was designed into a pcDNA3.1(+) backbone. Each of the ER-phagy receptors examined were tagged with mRFP-eGFP at either the N or C terminus (**Supplementary Table S6**) depending on previously published construct designs into a pcDNA3.1(+) backbone. AC16 cells (Millipore) at passage 7 or HeLa cells (ATCC) at passage 3 were plated onto a 6-well plate and cultured as described previously. Once cells were about 80% confluent, cells were transfected with 2.5 µg of the ER-phagy reporter or 5 µg of the ER-phagy receptors using lipofectamine 3000 (Thermo Scientific) according to kit instructions and as described above. The next day, transfected cells were passaged using trypsinization and plated on a 96-well plate at a density of 8000 cells for the ER-phagy reporter or 15000 for the ER-phagy receptors. Eight wells per timepoint were treated with either a DMSO vehicle control or thapsigargin and incubated for indicated timepoints. For the experiments using ubiquitin-like modifier activating enzyme 5 (UBA5)-in-1 (MCE), UBA5-in-1 was added at the indicated concentrations 2 hours before fixation. At the end of the incubation period, the cells were fixed with 4% PFA in PBS (Thermo Scientific) for 15 min at room temperature, washed 3 times with PBS for 5 mins each wash, labeled with NucBlue Fixed Cell Stain ReadyProbes reagent (Thermo Scientific), and washed 1 time with PBS before the addition of 0.1% sodium azide in PBS in each well. The cells were stored at 4°C until further processing. Imaging was performed using the Cell Insight CX7 LZR Pro High Content Screening Platform (Thermo Scientific) using the preprogrammed colocalization assay. The program reports the correlation coefficient between the fluorophores specified. These results were further processed in R and screened for the 90 cells with the most average nuclear size at each timepoint/condition. The results were tested for normality using a Shapiro-Wilk test then plotted with statistics using Wilcoxon rank sum test within timepoints.

### UFM1 immunoblot

AC16 cells (Millipore) at passage 7 were plated for three replicates and two wells per replicate for a 24-hour DMSO treated condition, a 24-hour 1 µM thapsigargin treated condition, and three UBA5-in-1 (MCE) concentrations added in conjunction to 24 hour 1 µM thapsigargin. After reaching about 80% confluency, the conditions were started, and UBA5-in-1 (MCE) was added 2 hours before harvesting. Cells were harvested with trypsin, pelleted at 300 ×g for 3 min and washed with 1 mL PBS by inverting 2x. Pellets were snap frozen using liquid nitrogen and stored at –80°C until further processing.

Pellets were resuspended, sonicated, and protein concentration was measured as described previously. 20 µg per sample was prepared for a final concentration of 30 µL with 4× Laemmli Sample Buffer (BioRad) and 10× NuPAGE Sample Reducing Agent (Thermo Scientific). Samples were loaded onto Mini PROTEAN TGX Stain Free Gels (BioRad) along with Precision Plus Protein Dual Color Standards (BioRad) and separated by electrophoresis using a constant 200 V for 30 min in Tris/Glycine/SDS Buffer (BioRad). The gels were transferred to a Trans-Blot Turbo Mini 0.2 µm PVDF Transfer Pack (BioRad) using a Turbo Trans Blot instrument (BioRad).

Membranes were blocked using 5% (w/v) Blotting Grade Blocker (BioRad) in 1x Tris-Buffered Saline (Fisher Scientific) with 0.1% (w/v) Tween (Millipore) for 1 hour mixing at room temperature and subsequently washed 3 times with 1× Tris-Buffered Saline (Fisher Scientific) with 0.1% (w/v) Tween (Millipore) for 5 mins mixing at room temperature. Membranes were then incubated with 1:1000 Rabbit anti-UFM1 (CST) in 1x Tris-Buffered Saline (Fisher Scientific) with 0.1% (w/v) Tween (Millipore) overnight mixing at 4 °C and subsequently washed 3 times with 1× Tris-Buffered Saline (Fisher Scientific) with 0.1% (w/v) Tween (Millipore) for 5 mins mixing at room temperature. This was followed by incubation with 1:1000 anti-rabbit IgG HRP conjugated (CST) in 1× Tris-Buffered Saline (Fisher Scientific) for 1 hour mixing at room temperature and 3 washes with 1× Tris-Buffered Saline (Fisher Scientific) for 5 mins mixing at room temperature. Membranes were imaged using SuperSignal West Pico PLUS Chemiluminescent Substrate (Thermo Scientific) on a ChemiDoc Imaging System (BioRad) using the chemiluminescent setting for 180 sec to 540 sec exposures with 4 images taken.

Antibodies were stripped off using Restore PLUS Western Blot Stripping Buffer (Thermo Scientific) and the membrane was rinsed with Tris-Buffered Saline (Fisher Scientific) for 5 mins mixing at room temperature. The membrane was sprayed with methanol and incubated with Ponceau S Solution 0.1% (BIOTUM) for 10 mins mixing at room temperature. Membrane was transferred to Tris-Buffered Saline (Fisher Scientific) and imaged on ChemiDoc Imaging System (BioRad) using the Ponceau setting for total protein detection.

### Cell survival assay

For ER-phagy constructs, AC16 cells (Millipore) were plated onto a 6-well plate. Once cells were about 80% confluent, cells were transfected with 2.5 µg of the ER-phagy reporter or 5 µg of the ER-phagy receptors and split onto a 96-well plate at a density of 10000 cells per well as previously described. For non-transfected experiments, AC16 cells (Millipore) at passage 8 or HeLa (ATCC) cells at passage 4 were plated onto a 96-well plate at a density of 5000 cells per well and cultured as previously described. For testing of UBA5-in-1 on viability, the cells were exposed to either DMSO or 1 µM thapsigargin for 24 hours and UBA5-in-1 was added at indicated concentrations 2 hours before the addition of resazurin (CST). Resazurin (CST) was thawed at 37 °C and added at a ratio of 1:10 to each well before incubation at 37 °C for 5 hours. Resazurin conversion to resorufin indicating metabolic activity was read using the Spectral Max iD5 (Molecular Devices) with either fluorescence (emission and excitation at 570nm/620nm) or normalized absorbance for wavelengths 570nm/620nm^31^. Experiments performed on transfected cells were examined using the cyQUANT XTT assay (Invitrogen) at 450nm absorbance using the Spectral Max iD5 (Molecular Devices) after 4 hours of incubation at 37 °C.

### UFM1 pulldown

AC16 cells (Millipore) passage 7 were passaged onto four 150mm cell culture dishes and grown to 90% confluency as previously described. Two plates were treated with 1 µM thapsigargin for 24 hours before harvesting. Briefly, media was removed, plates were rinsed with PBS 2× and incubated on ice with 1.2 mL of 10x Cell Lysis Buffer (CST) diluted to 1x for 5 mins before scraping. The harvested cells were combined and sonicated in a Bioruptor sonicator with settings 10× 30 s on 30 s off at 4°C, insoluble debris were removed using centrifugation at 14,000 x g at 4°C for 5 mins, and protein concentration was determined using a Pierce BCA protein assay kit (Thermo Scientific). 100 µL of Protein A magnetic beads (CST) was aliquoted, separated from storage solution, and rinsed 2 times with diluted 10x Cell Lysis Buffer (CST) using a magnetic rack. To remove background, 1 mg of sample was adjusted to a final volume of 1 mL and incubated with the beads for 20 mins rotating at 22°C. The supernatant was incubated with 1:50 antibody to protein Rabbit anti-UFM1 (CST) overnight rotating at 4°C. 100 µL of Protein A magnetic beads (CST) was prepared as previously described using 10x Cell Lysis Buffer (CST) diluted in LC-MS grade water. To pull down the UFM1 antibody, the sample was incubated with the prepared beads for 20 mins rotating at 22°C. The beads were collected and washed using 1 mL of 10x Cell Lysis Buffer (CST) diluted in LC-MS grade water 5 times and 1 mL 20mM Tris-HCl pH 7.5 5 times on a magnetic rack. The beads were resuspended in 30 µL of 100 ng/µl of mass spectrometry grade trypsin (Promega) in 20mM Tris-HCl pH7.5 and incubated overnight at 37°C. The sample was dried in a speed-vac and reconstituted in 10 µL of pH 2 MS solvent A (0.1% formic acid) before triplicate analysis with LC-MS/MS on a Thermo Exploris 480 orbitrap mass spectrometer coupled to an LC with electrospray ionization source. Peptides were separated with a PepMap RSLC C18 column 75 µm x 15 cm, 3 µm particle size (Thermo Scientific) using a 117-minute gradient from 0 to 99% pH 2 MS solvent B (0.1% formic acid in 80% LC-MS grade acetonitrile) with a flow rate of 200nL/min. The gradient of solvent B increased from 2–20% for 76 minutes, 20–30% for 27.5 minutes, 30–60% for 11.5 minutes, 60–99% for 0.9 minutes, and held at 99% for 1 minute. The mass spectrometer operated in positive mode using a source voltage of 2000 V with MS1 scans acquired from 400–900 m/z at 45,000 resolution in profile mode using a maximum injection time set to auto with a normalized AGC target of 300% and RF lens of 50%. Peptide precursor ions were selected following data-independent acquisition with 6 m/z isolation windows from 400–900 m/z with optimized placement. The maximum ion accumulation time was set to auto and the normalized AGC target set to 1000% and fragmented by HCD at 29 NCE. MS2 scans were acquired with a fixed first mass set to 120 m/z at 22,500 resolution in profile mode with loop control set to all. Raw spectra were converted using DIA-NN 2.2.0 with missed cleavages set to 1, variable modifications set to 2, peptide length range of 7-45, precursor charge range of 2-5, precursor m/z range set to 400-900, and fragment ion m/z range set to 120-1870 using the UniProt Swiss-Prot human canonical and isoform protein sequence database (retrieved 2025-08-29). The output was filtered by proteotypic = 1, protein Q value < 0.01, and global protein group Q value < 0.01 before comparing pulled down proteins to the BioGRID database interactome (93) of each indicated ER-phagy receptor.

### Quantification and statistical analysis

Quantification and statistical analysis are described in the analysis of each respective technique above. Unless otherwise specified, protein identification requires decoy database assisted calculation of 1% peptide and protein FDR. Protein differential localization requires BANDLE 99% probability. Differential synthesis and clearance require limma P value < 0.05. Skewed datasets were assessed with a Wilcoxon rank sum test with significance cutoff values as 0.05 for a single star, 0.01 for two stars, and each additional star representing another magnitude of significance. Data sets that did not show obvious skewness were assessed using t-tests with identical significance cut off values. For the Seahorse assays, multiple stages were compared at each timepoint using an ANOVA with pairwise comparisons adjusted using the Holm method. Assessments dependent on Mahalanobis distance relied on chi-squared cut offs of 90-95%.

## Supplementary Information

**Supplementary Figure S1.**
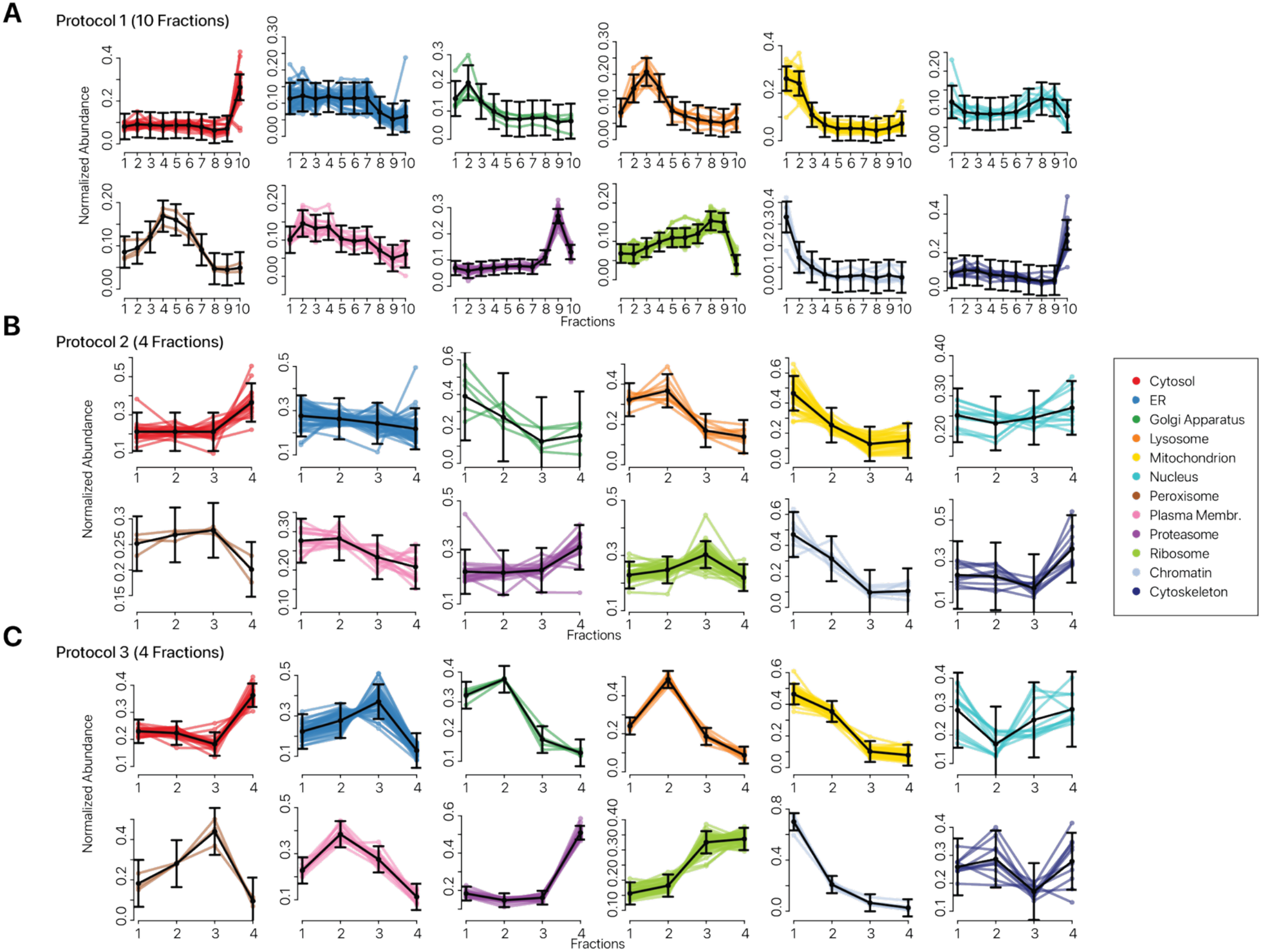
Fraction profiles of reduced fractionation protocols. (A) Distribution patterns of normalized abundance (y-axis) from TMT channel intensities for each subcellular marker over the ultracentrifugation fractions (x-axis) for standard 10-fraction SPLAT/LOPIT-DC protocol (Protocol 1). Black lines depict the average intensity at each fraction for the markers of that compartment with error bars for the standard deviation. Data points are from technical triplicate experiments. (B) As in A, but for the reduced 4-fraction Protocol 2. (C) As in A, but for the reduced 4-fraction Protocol 3.

**Supplementary Figure S2.**
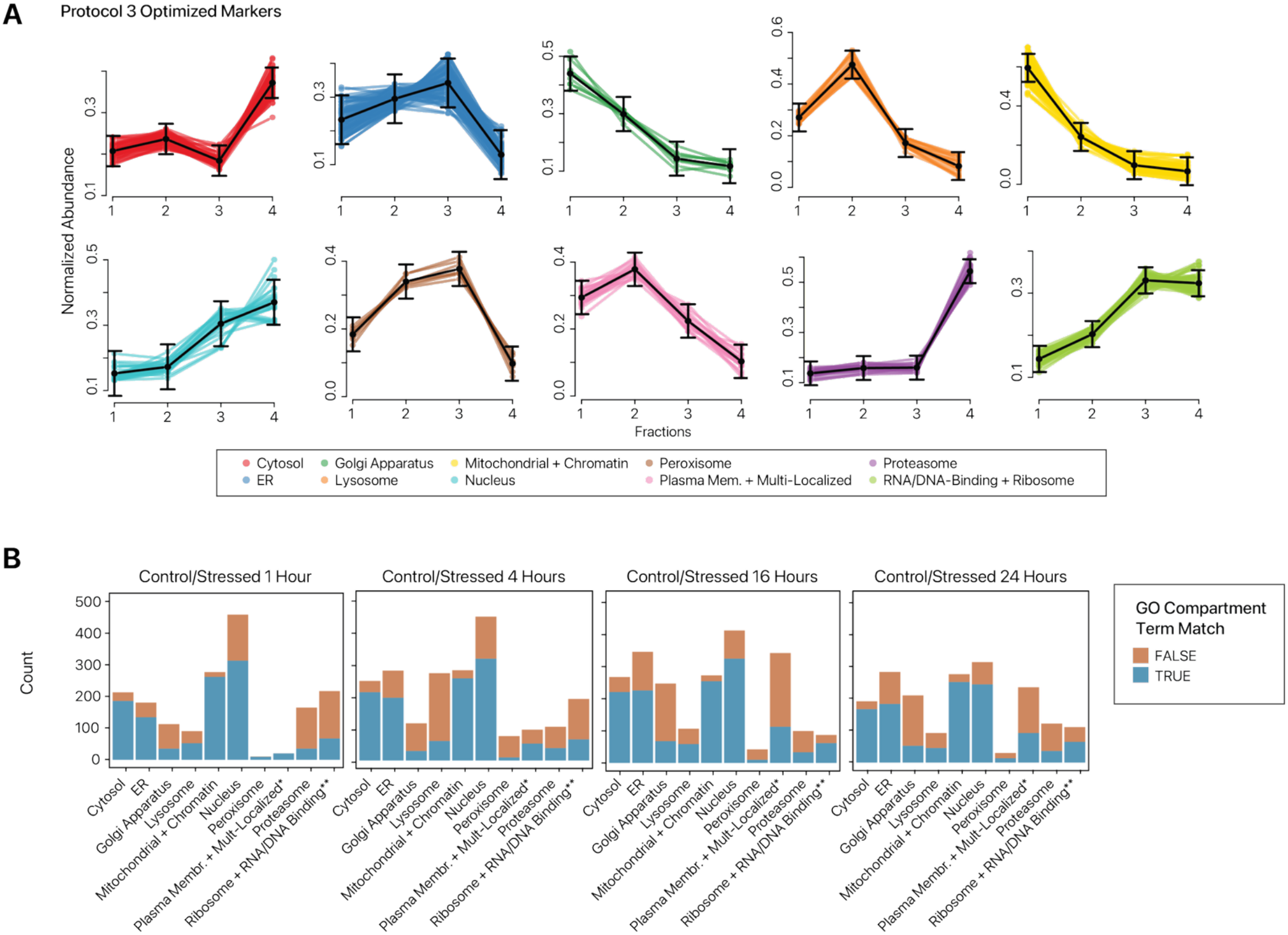
Fraction profiles of optimized localization markers in SPLAT-TR. (A) Distribution patterns of normalized abundance (y-axis) calculated from TMT channel intensities for each subcellular marker over the ultracentrifugation fractions (x-axis) for the SPLAT-TR markers. Black lines depict the average intensity at each fraction for the markers of that compartment with error bars for the standard deviation. Data are from the first replicate of the 1hr control condition. (B) Bar charts comparing the BANDLE assigned subcellular localization of each timepoint with the GO Cellular Compartment term of the proteins. For each timepoint, replicates from both the control and stressed conditions are trained together, and the control cell localizations are graphed. For the Plasma Membrane + Multi-Localized compartment, the experiential assignments are compared to plasma membrane annotations only. For the “Ribosome + RNA/DNA Binding” compartment, the experiential assignments are compared to ribosome annotations only.

**Supplementary Figure S3.**
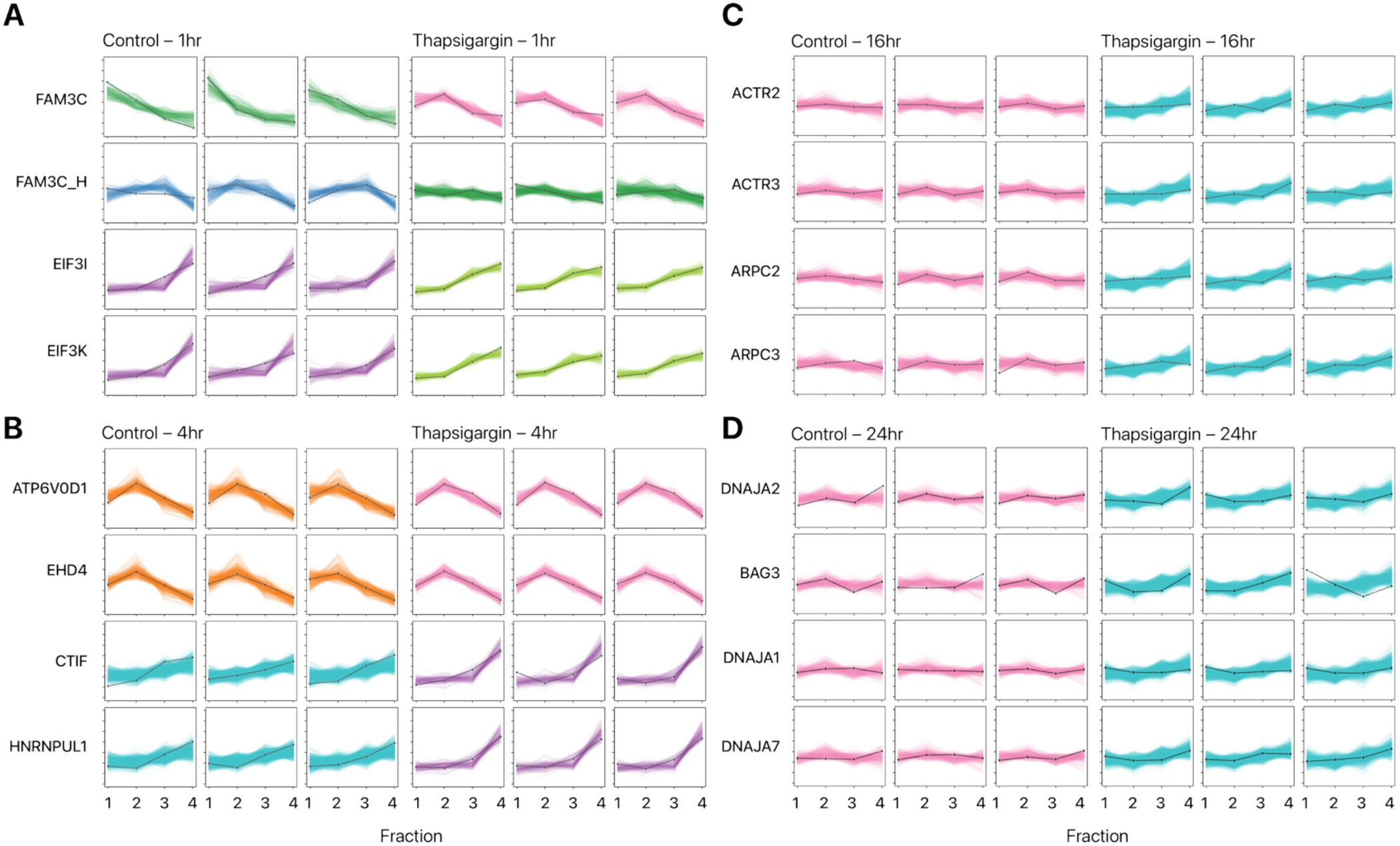
Protein distribution profiles for sequential differential localization events. (A) Protein distribution profiles of a subset of the proteins predicted to re-locate in Figure 2 for the 1-hour stressed timepoint. Lines are the normalized abundance (y-axis) ranging from 0 on the first dash to 0.7 on the last dash over each fraction (x-axis), 1 through 4 with a dash for each fraction. Line color represents predicted subcellular localization. The black line is the distribution pattern for protein indicated, and the colored lines are the distribution patterns for all the markers of that subcellular region. The color scheme follows that of Figure 2. (B) As in A, but for the 4-hour stressed timepoint. (C) As in A, but for the 16-hour stressed timepoint. (D) As in A, but for the 24-hour stressed timepoint.

**Supplementary Figure S4.**
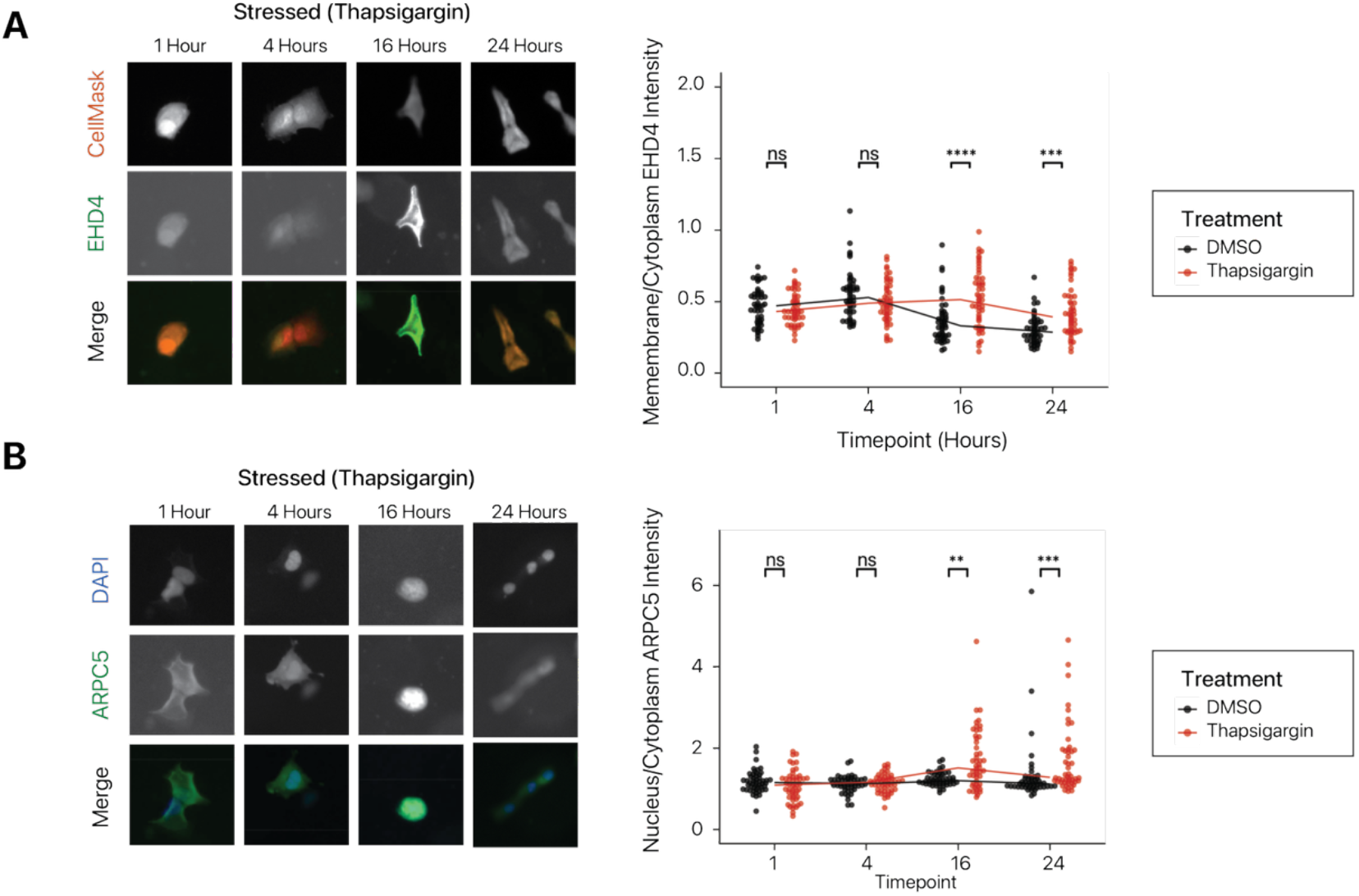
Validation experiments for additional differential localization proteins. (A) Left: High-content imaging of SNAP-tagged constructs for EHD4 in stressed AC16 cells at 1 hour, 4 hours, 16 hours, and 24 hours. Deep read: CellMask for whole cell label; Oregon Green: SNAP ligand. Right: Analysis of the intensity of the Oregon Green SNAP ligand at the membrane over the intensity in the cytoplasm (y-axis) at each timepoint (x-axis). Each timepoint contains 50 cells screened for the highest expression for the protein of interest of the cells imaged at that timepoint. A Shapiro-Wilk test indicated the data deviated from normality. Significance tests were performed using a Wilcoxon Rank Sum Test comparing the control and thapsigargin treated at each timepoint. Solid line indicates the median of each condition. (B) Left: High-content imaging of SNAP-tagged constructs for ARPC5 in stressed AC16 cells at 1 hour, 4 hours, 16 hours, and 24 hours. Blue: nucleus (DAPI); Oregon Green: SNAP ligand. Right: Analysis of the intensity of the Oregon Green SNAP ligand at the nucleus (overlapping DAPI) over the intensity in the cytoplasm (y-axis) at each timepoint (x-axis). Each timepoint contains 50 cells screened for most average nuclear size of the cells imaged at that timepoint. A Shapiro-Wilk test indicated the data deviated from normality. Significance tests were performed using a Wilcoxon Rank Sum Test comparing the control and thapsigargin treated at each timepoint. Solid line indicates the median of each condition.

**Supplementary Figure S5.**
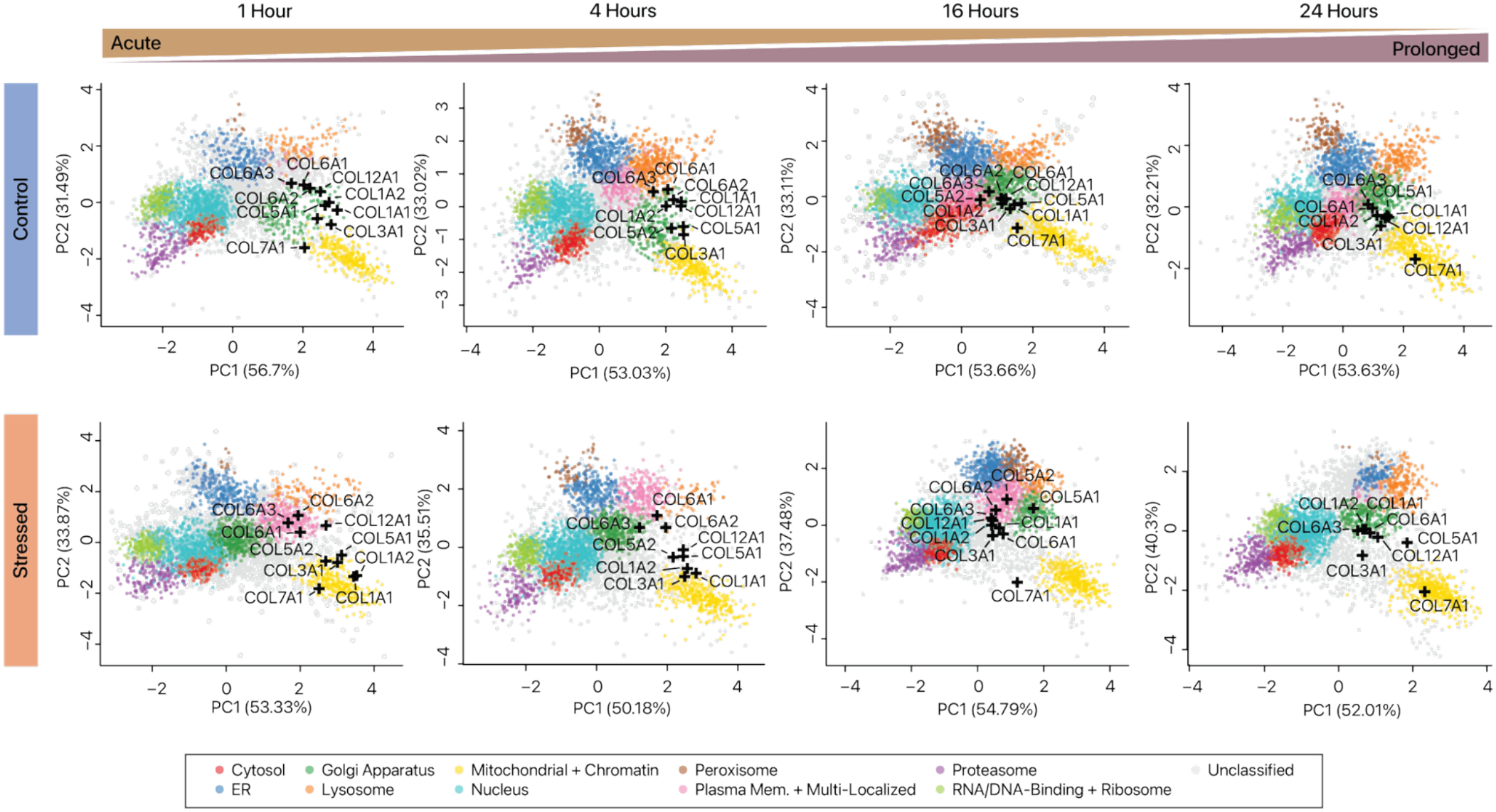
Differential localization of collagen protein chains under ER stress. Spatial maps are shown with the subcellular locations (black cross) of unlabeled collagen chains which were predicted to re-locate at any timepoint. Top panels show the control localizations, and bottom panels show the thapsigargin treated localizations. PCAs portrayed are an average of the three replicates collected. Legend for color scheme is shown at the bottom.

**Supplementary Figure S6.**
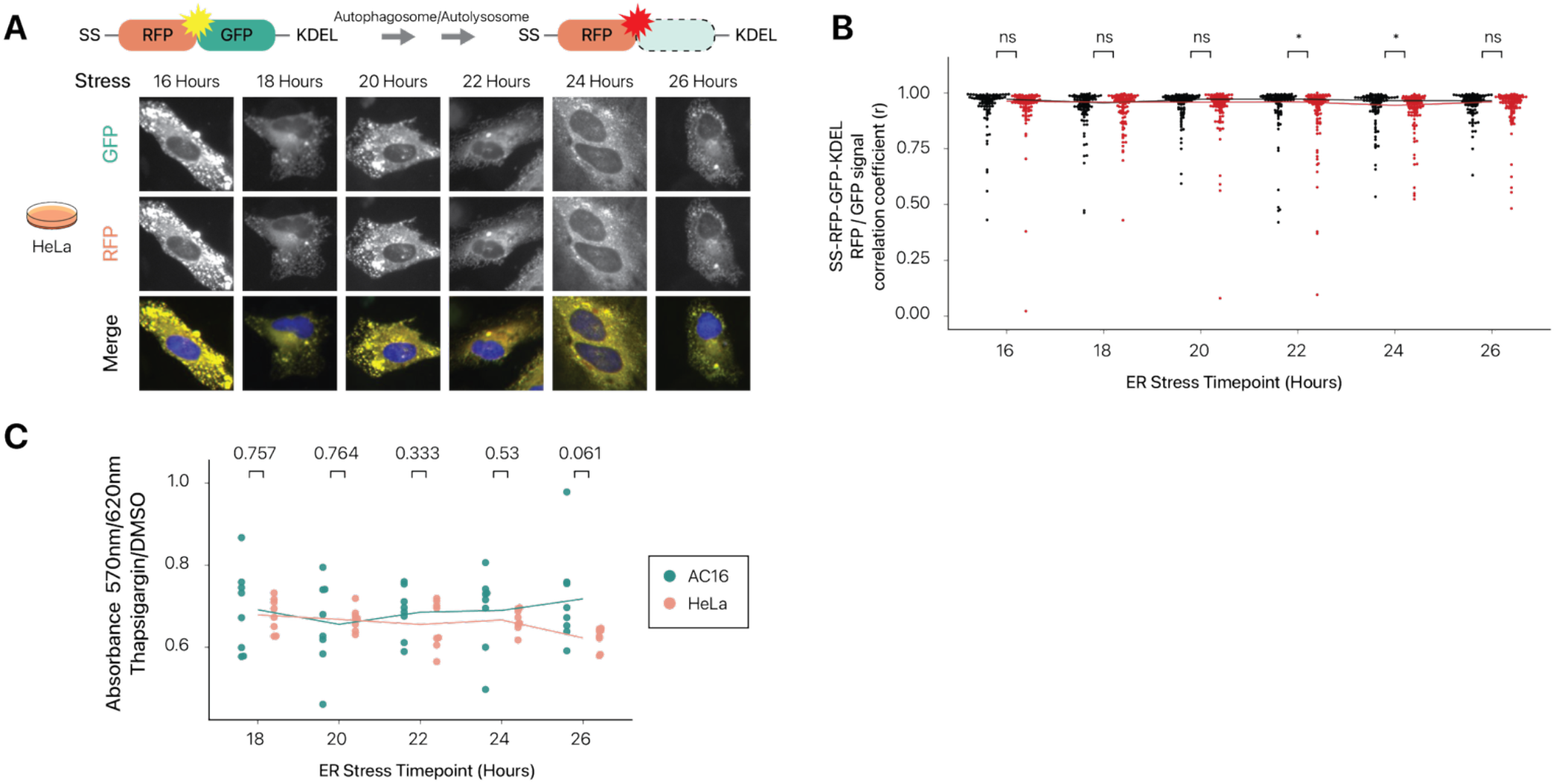
Cell line dependency on ER-phagy correlates with sensitivity to ER stress. (A) High content imaging of a 1 µM thapsigargin time course on HeLa cells using ER-phagy assay. (B) Plots of the analysis of the high content screening results for the correlation coefficient between the RFP and GFP signal (y-axis) over a time course of 1 µM thapsigargin treatment (x-axis). Each timepoint contains 90 cells screened for most average nuclear size of the cells imaged at that timepoint. DMSO treated samples (black) and thapsigargin treated samples (red) were compared at each timepoint using a Wilcoxon Rank Sum Test after a Shapiro-Wilk test indicated the data deviated from normality. Solid line indicates the mean of each condition. (C) Plot of a resazurin fluorescent viability assay with the ratio of the absorbance at 570nm over 620nm normalized to the mean of the DMSO treated condition (y-axis) for multiple timepoints (x-axis) in AC16 and HeLa cells. Statistics were performed using a t-test between indicated timepoints. Each timepoint was performed with eight replicates. Solid line indicates the mean of the timepoint.

**Supplementary Table S1.** Differential ultracentrifugation protocols.

|  |  |  |
| --- | --- | --- |
| **Protocol 1** |  |  |
| **Sample** | **Speed** | **Time** |
| Pellet 1 | 1,000 × g | 10 min |
| Pellet 2# | 3,000 × g | 10 min |
| Pellet 3# | 5,000 × g | 10 min |
| Pellet 4 | 9,000 × g | 15 min |
| Pellet 5# | 12,000 × g | 15 min |
| Pellet 6 | 15,000 × g | 15 min |
| Pellet 7# | 30,000 × g | 20 min |
| Pellet 8 | 79,000 × g | 43 min |
| Pellet 9 | 120,000 × g | 45 min |
| Supernatant 10# |  |  |
| **#: Kept in Protocol 2** |  |  |
| **Protocol 3** |  |  |
| **Sample** | **Speed** | **Time** |
| Pellet 1 | 3,000 × g | 10 min |
| Pellet 2 | 9,000 × g | 15 min |
| Pellet 3 | 30,000 × g | 20 min |
| Pellet 4 | 120,000 × g | 45 min |
| Supernatant 5 |  |  |

**Supplementary Table S2.** Fraction association of TMT channels in control and ER stressed AC16. Abbreviations: C = Control; T = Thapsigargin; F = Fraction.

| **TMT Label** | **Replicate 1** | **Replicate 1** | **Replicate 2** | **Replicate 2** | **Replicate 3** | **Replicate 3** |
| --- | --- | --- | --- | --- | --- | --- |
| 126 | C 1hr F1 | C 24hr F1 | C 1hr F4 | C 24hr F3 | T 1hr F1 | C 4hr F3 |
| 127N | C 1hr F3 | T 4hr F1 | C 16hr F1 | T 24hr F3 | T 1hr F2 | T 24hr F1 |
| 127C | C 1hr F4 | T 24hr F4 | T 1hr F4 | C 4hr F1 | T 1hr F4 | T 24hr F4 |
| 128N | T 16hr F2 | T 4hr F2 | C 16hr F4 | T 4hr F2 | C 1hr F4 | C 24hr F3 |
| 128C | C 16hr F3 | T 4hr F4 | T 1hr F2 | C 24hr F1 | C 1hr F2 | C 4hr F1 |
| 129N | C 16hr F1 | C 24hr F3 | C 16hr F3 | C 24hr F2 | C 16hr F3 | T 4hr F2 |
| 129C | T 1hr F2 | T 24hr F1 | T 16hr F4 | T 24hr F2 | T 16hr F1 | T 24hr F3 |
| 130N | T 16hr F4 | C 24hr F2 | T 1hr F3 | T 4hr F3 | C 16hr F1 | T 4hr F4 |
| 130C | T 1hr F4 | C 4hr F4 | C 16hr F2 | T 4hr F4 | C 1hr F3 | T 4hr F3 |
| 131N | T 16hr F3 | T 4hr F3 | C 1hr F1 | C 4hr F2 | T 16hr F4 | C 4hr F4 |
| 131C | C 16hr F2 | C 4hr F2 | T 16hr F2 | T 4hr F1 | C 1hr F1 | T 4hr F1 |
| 132N | T 1hr F1 | T 24hr F3 | T 1hr F1 | T 24hr F4 | T 16hr F3 | C 4hr F2 |
| 132C | C 1hr F2 | T 24hr F2 | C 1hr F2 | C 4hr F3 | T 1hr F3 | T 24hr F2 |
| 133N | C 16hr F4 | C 4hr F1 | T 16hr F1 | T 24hr F1 | T 16hr F2 | C 24hr F2 |
| 133C | T 1hr F3 | C 4hr F3 | T 16hr F3 | C 4hr F4 | C 16hr F2 | C 24hr F1 |
| 134N | T 16hr F1 | C 24hr F4 | C 1hr F3 | C 24hr F4 | C 16hr F4 | C 24hr F4 |

**Supplementary Table S3.** Isotopic contaminant matrix of TMTpro-16plex lot XL348283.

| **TMT Label** | **-2x 13C** | **-13C; -15N** | **-13C** | **-15N** | **+15N** | **+13C** | **+15N; +13C** | **+2x 13C** |
| --- | --- | --- | --- | --- | --- | --- | --- | --- |
| 126 | N/A% | N/A% | N/A% | N/A% | 0.31% | 9.09%<br>(127C) | 0.02% | 0.32% |
| 127N | N/A% | N/A% | N/A% | 0.78%<br>(126) | N/A% | 9.41%<br>(128N) | N/A% | 0.33% |
| 127C | N/A% | N/A% | 0.84%<br>(126) | N/A% | 0.23% | 8.40%<br>(128C) | 0.02% | 0.27% |
| 128N | N/A% | 0.00% | 0.82%<br>(127N) | 0.65% | N/A% | 8.13%<br>(129N) | N/A% | 0.26% |
| 128C | 0.00% | N/A% | 1.44%<br>(127C) | N/A% | 0.34% | 6.26%<br>(129C) | 0.00% | 0.17% |
| 129N | 0.00% | 0.14% | 1.30%<br>(128N) | 0.89% | N/A% | 7.52%<br>(130N) | N/A% | 0.12% |
| 129C | 0.13% | N/A% | 2.59%<br>(128C) | N/A% | 0.32% | 6.07%<br>(130C) | 0.01% | 0.09% |
| 130N | 0.13% | 0.00% | 2.41%<br>(129N) | 0.27% | N/A% | 5.58%<br>(131N) | N/A% | 0.10% |
| 130C | 0.25% | N/A% | 3.22%<br>(129C) | N/A% | 0.28% | 5.06%<br>(131C) | 0.00% | 0.06% |
| 131N | 0.03% | 0.00% | 2.78%<br>(130N) | 0.63% | N/A% | 4.57%<br>(132N) | N/A% | 0.12% |
| 131C | 0.08% | N/A% | 3.94%<br>(130C) | N/A% | 0.45% | 3.33%<br>(132C) | 0.00% | 0.02% |
| 132N | 0.07% | 0.01% | 3.14%<br>(131N) | 0.73% | N/A% | 3.40%<br>(133N) | N/A% | 0.03% |
| 132C | 0.08% | N/A% | 3.65%<br>(131C) | N/A% | 0.50% | 1.97%<br>(133C) | 0.00% | 0.00% |
| 133N | 0.15% | 0.01% | 3.58%<br>(132N) | 0.72% | N/A% | 1.80%<br>(134N) | N/A% | 0.00% |
| 133C | 0.22% | N/A% | 4.96%<br>(132C) | N/A% | 0.34% | 1.03%<br>(134C) | 0.00% | N/A% |
| 134N | 0.30% | 0.03% | 5.49%<br>(133N) | 0.62% | N/A% | 1.14%<br>(135N) | N/A% | N/A% |

**Supplementary Table S4.** Isotopic contaminant matrix of TMTpro-16plex lot YB367250.

| **TMT Label** | **-2x 13C** | **-13C; -15N** | **-13C** | **-15N** | **+15N** | **+13C** | **+15N; +13C** | **+2x 13C** |
| --- | --- | --- | --- | --- | --- | --- | --- | --- |
| 126 | N/A | N/A | N/A | 0.57% (126) | 0.31% | 9.09% (127C) | 0.02% | 0.32% |
| 127N | N/A | N/A | 0.84% (126) | N/A | N/A | 9.79% (128N) | N/A | 0.33% |
| 127C | N/A | 0.00% | 0.82% (127N) | 0.65% | 0.23% | 8.40% (128C) | 0.02% | 0.27% |
| 128N | 0.00% | N/A | 1.44% (127C) | N/A | N/A | 8.13% (129N) | N/A | 0.26% |
| 128C | 0.00% | 0.14% | 1.30% (128N) | 0.89% | 0.34% | 6.26% (129C) | 0.00% | 0.17% |
| 129N | 0.13% | N/A | 2.59% (128C) | N/A | N/A | 7.52% (130N) | N/A | 0.12% |
| 129C | 0.13% | 0.00% | 2.41% (129N) | 0.27% | 0.32% | 6.07% (130C) | 0.01% | 0.09% |
| 130N | 0.25% | N/A | 3.22% (129C) | N/A | N/A | 5.58% (131N) | N/A | 0.10% |
| 130C | 0.03% | 0.00% | 2.78% (130N) | 0.63% | 0.28% | 5.06% (131C) | 0.00% | 0.06% |
| 131N | 0.08% | N/A | 3.94% (130C) | N/A | N/A | 4.57% (132N) | N/A | 0.12% |
| 131C | 0.07% | 0.01% | 3.14% (131N) | 0.73% | 0.45% | 3.33% (132C) | 0.00% | 0.02% |
| 132N | 0.08% | N/A | 3.65% (131C) | N/A | N/A | 3.40% (133N) | N/A | 0.03% |
| 132C | 0.15% | 0.01% | 3.58% (132N) | 0.72% | 0.50% | 1.97% (133C) | 0.00% | 0.00% |
| 133N | 0.22% | N/A | 4.96% (132C) | N/A | N/A | 1.80% (134N) | N/A | 0.00% |
| 133C | 0.30% | 0.03% | 5.49% (133N) | 0.62% | 0.34% | 1.03% (134C) | 0.00% | N/A |
| 134N | N/A | N/A | N/A | 0.57% (126) | N/A | 1.14% (135N) | N/A | N/A |

**Supplementary Table S5.** Fraction association of TMTpro channels in protein degradation inhibition experiments. Numbers signify duration of thapsigargin exposure. T = Thapsigargin; M = MG132; B = Bafilomycin A1; R = Replicate

| **TMT Label** | **Sample** |
| --- | --- |
| 126 | M6 R1 |
| 127N | T24 R3 |
| 127C | B6 R1 |
| 128N | C24 R2 |
| 128C | M6 R3 |
| 129N | M6 R2 |
| 129C | T6 R3 |
| 130N | T24 R1 |
| 130C | T6 R1 |
| 131N | T6 R2 |
| 131C | B24 R1 |
| 132N | B6 R3 |
| 132C | B24 R3 |
| 133N | B24 R2 |
| 133C | B6 R2 |
| 134N | M6 R1 |

**Supplementary Table S6.** Construct outline of ER-phagy receptor-tagged RFP-GFP reporters. MCS = Multiple Cloning Site; SS = BIP signal sequence

| Construct |
| --- |
| pcDNA3.1(+)-SS-mRFP-MCS-eGFP-KDEL |
| pcDNA3.1(+)-CYB5R3-linker-mRFP-MCS-eGFP |
| pcDNA3.1(+)-TEX264-linker-mRFP-MCS-eGFP |
| pcDNA3.1(+)-mRFP-MCS-eGFP-linker-CCPG1 |
| pcDNA3.1(+)-mRFP-MCS-eGFP-linker-FAM134A |
| pcDNA3.1(+)-mRFP-MCS-eGFP-linker-FAM134B |
| pcDNA3.1(+)-mRFP-MCS-eGFP-linker-FAM134C |

